# HIV disrupts CD34^+^ progenitors involved in T-cell differentiation

**DOI:** 10.1101/142539

**Authors:** Tetsuo Tsukamoto

## Abstract

HIV-1 causes the loss of CD4^+^ T cells via depletion or impairment of their production. The latter involves infection of thymocytes, but the involvement of other cells including haematopoietic CD34^+^ cells remains unclear even though HIV-positive patients frequently manifest myelosuppression. This study utilised the OP9-DL1 coculture system, which supports in vitro T-lineage differentiation of human haematopoietic stem/progenitor cells. Cord-derived CD34^+^ cells were infected with CXCR4-tropic HIV-1_NL4-3_ and cocultured. HIV-infected cocultures exhibited sustained viral replication for 5 weeks, as well as reduced CD4^+^ T-cell growth at weeks 3–5. It was further revealed that CD34^+^CD7^+^CXCR4^+^ cells can be quickly depleted as early as in 1 week after infection of the subset, and this was accompanied by the emergence of CD34^+^CD7^+^CD4^+^ cells. These results indicate that CXCR4-tropic HIV-1 strains may disrupt CD34^+^CD7^+^ lymphoid progenitor cell pools, presumably leading to impaired T-cell production potential.

## Introduction

Human immunodeficiency virus (HIV) infection is associated with haematological changes ^1^. Antiretroviral therapy is effective in controlling viremia and treating acquired immunodeficiency syndrome (AIDS). However, some patients do not experience sufficient T-cell immune restoration despite being aviremic during treatment ^2^. HIV-infected patients may display decreased thymopoiesis ^3^. Thymic dysfunction occurs during HIV infection, and it is associated with rapid progression in infants with prenatal HIV infection ^4^. A previous report tested coculture of CD34^+^ and fetal thymic epithelial cells in the presence or absence of HIV-1, observing that infection led to the inhibition of thymocyte maturation at early stages (CD44+CD25-CD3-) ^5^. Pancytopenia may occur as a result of bone marrow abnormalities in HIV-infected individuals ^6,7^. A previous study revealed depletion of CD34^+^CD4^+^ cells in bone marrow from HIV-infected patients. However, the study failed to uncover evidence of HIV infection of these cells ^8^.

The thymopoiesis environment can be mimicked in vitro using the OP9-DL1 or OP9-DL4 cell line. These cell lines were derived from the OP9 mouse stromal cell line via transduction with a notch ligand called delta-like 1 (DL1) or DL4 ^9,10^. The ability of OP9-DL1 cells to support thymopoiesis in vitro was first demonstrated in coculture with mouse cells ^9^. The cell line is also known to support the differentiation of human CD34^+^ haematopoietic stem/progenitor cells (HSPCs) into thymocytes and T cells ^11^. There is evidence that OP9-DL1 cells produce stromal derived factor-1 (SDF-1, also known as CXCL12), a ligand for CXCR4 ^12^. Although OP9-DL4 cells can induce differentiation into both specific myeloid cells and T-lineage cells, OP9-DL1 cells only permit the differentiation of T-lineage cells while inhibiting B cells and myeloid cells ^13^. Thus, the OP9-DL1 coculture system is used to investigate events associated with T-lineage differentiation.

The chemokine receptors CXCR4 and CCR5 are common coreceptors for HIV-1 ^14^. Control of CCR5-tropic strains of HIV-1 is usually considered a better correlate of good clinical outcomes than control of CXCR4-tropic HIV-1 strains ^15^. This is due to the fact that memory CD4^+^ T cells express higher levels of CCR5, thus making them susceptible to CCR5-tropic HIV infection and subsequent depletion ^16^. Consequently, CCR5 may appear to be more closely involved in the immunopathogenesis of HIV infection than CXCR4 ^17^. Conversely, the biological functions of CXCR4 have been well documented in fields such as developmental biology and haematology. CXCR4 interacts with SDF-1 and allows CXCR4-expressing cells to home to loci in which SDF-1 is highly expressed ^18^. SDF-1 and CXCR4 are essential in human stem cell homing and repopulation of the host with differentiated haematopoietic cells ^19,20^. SDF-1 is also produced by thymic epithelial cells, and it plays an important role in migration of immature progenitors in the thymus ^21^.

Therefore, it may be important to better understand the influence of CXCR4-tropic HIV-1 infection on CXCR4-associated biological events including haematopoiesis and T-cell development ^22^. A prior study sought to clarify the effect of CXCR4-tropic simian-human immunodeficiency virus infection on T-lineage cell production in the thymi of newborn rhesus macaques ^23^. However, it is not realistic to closely follow in vivo bone marrow/thymus events in HIV-infected individuals. Instead, humanised mouse models can be beneficial for this purpose ^24,25^. Moreover, an easy-to-use ex-vivo model may be helpful for closely monitoring the differentiation of HSPCs into T-lineage cells in the presence of HIV-1. The present study aimed to determine the in vitro fate of CD34^+^ progenitor cells and derivatives exposed to HIV-1 using the OP9-DL1 coculture system.

## Results

### Persistent HIV-1 infection observed in OP9-DL1 cocultures with cord-derived CD34^+^cells

Two separate in vitro experiments were performed in the study. The data for experiment 1 (long coculture) are shown in Figs. 1–3 and Extended Data Figs. 1–5. To follow the in vitro fate of HIV–pre-exposed CD34^+^ cells and derivatives for several weeks, primary human umbilical cord-derived CD34^+^ cells were infected with HIV-1. These cells were partially CD4^+^ and/or CXCR4^+^ before infection (Extended Data Fig. 1A). Cells were seeded in 48-well plates and exposed to 200 ng (p24) of HIV-1NL4-3. Following centrifugation and overnight incubation in the presence of HIV-1, the cells were cocultured with OP9-DL1 cells (Fig. 1A). After a week of coculture, intracellular HIV-1 p24 was detected in both CD34^+^ and CD34^-^ cells (Fig. 1B). HIV infection was further examined via magnetic bead separation of CD34^+^ and CD34^-^ cells followed by DNA extraction and detection of HIV-1 *gag* DNA using PCR (Extended Data Fig. 1B). The HIV-infected samples displayed persistent HIV-1 p24 expression in 0.1–1.7% of the total cocultured human cells (Fig. 1C, Extended Data Fig. 1C–G). A correlation was found between intracellular HIV-1 p24^+^ cell counts and supernatant HIV-1 p24 concentrations (Extended Fig. 1E).

**Figure 1.**
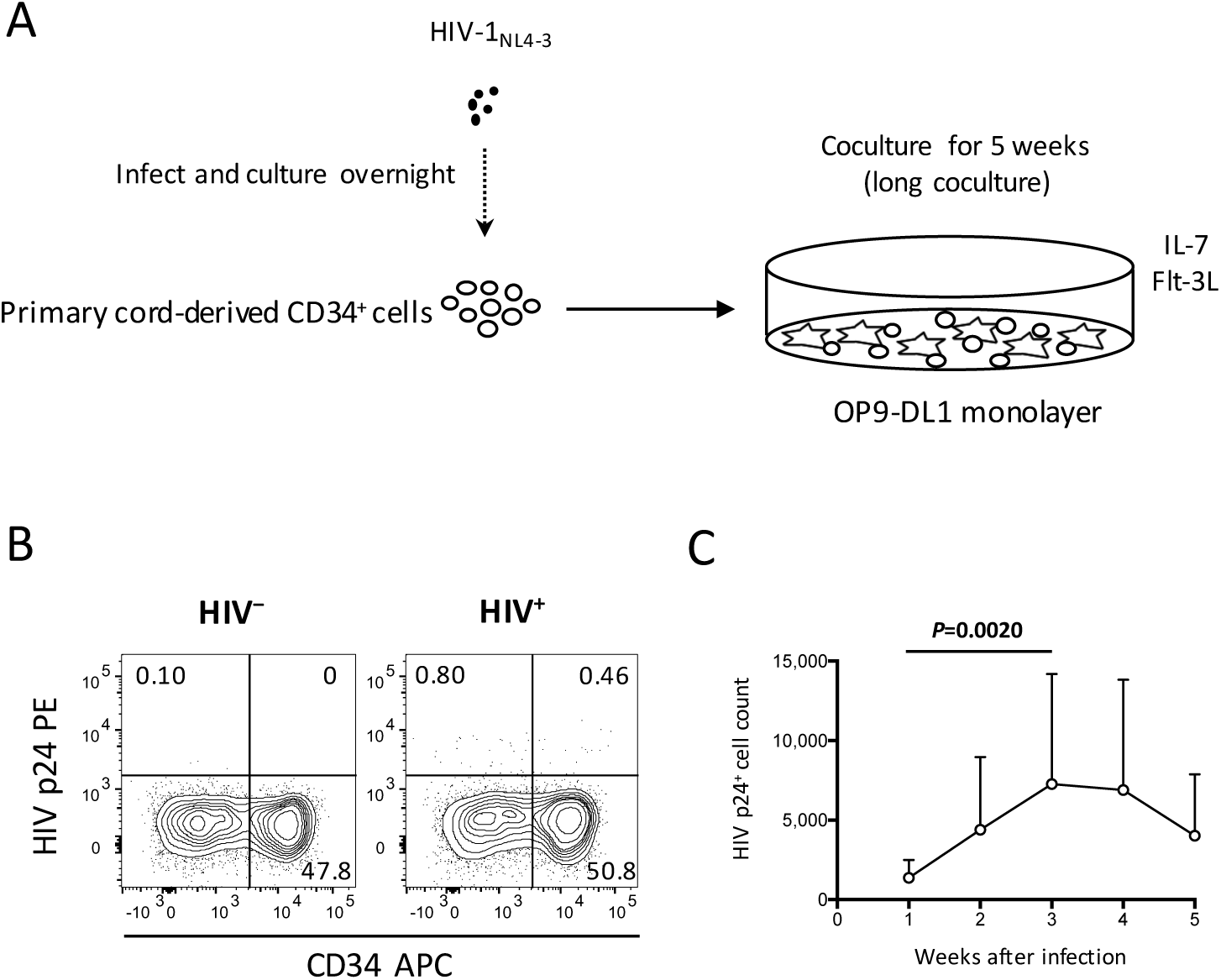
Primary umbilical cord-derived CD34^+^ cells are susceptible to HIV infection. Coculture of HIV-infected primary cord-derived CD34^+^ cells with OP9-DL1 cells resulted in persistent viral replication for 5 weeks (n = 12). (A) Schematic representation of experiment 1 (long coculture). Cord-derived CD34^+^ cells were infected with HIV-1_NL4-3_ and cocultured with OP9-DL1 cells for 5 weeks. (B) Intracellular HIV p24 expression was tested 1 week after infection. HIV p24^+^ cells were found in both the CD34^+^ and CD34^-^fractions of HIV-infected samples. Representative plots are shown. (C) HIV p24^+^ cell counts at weeks 1–5 post-infection (n = 12). Statistical analyses were performed using the Wilcoxon matched-pairs signed rank test.

### The dynamics of CD4^+^ cells in OP9-DL1 cocultures with HIV–pre-exposed CD34^+^ cells

The influence of HIV pre-exposure of CD34^+^ cells on post-coculture events was analysed by flow cytometry weekly until week 5. CD19^+^, CD20^+^ or CD33^+^ cells were not detected in the samples tested (data not shown). HIV-infected cocultures had significantly lower whole cell counts at weeks 3, 4 and 5 post-coculture than uninfected cocultures (Extended Data Fig. 2A), demonstrating reduced cell growth in the presence of HIV-1. Similarly, the CD4^+^CD8^+^ and CD4^+^CD8^+^CXCR4^+^ cell counts were significantly smaller in HIV^+^ cocultures than in HIV^-^ cocultures at week 3 (Extended Data Fig. 2B–D). When CD34^-^ cells were gated and analysed for CD7 and CD4 expression (Fig. 2A), the CD34^-^CD7^+^CD4^+^ cell counts were not significantly different between HIV-infected and uninfected samples (Fig. 2B). By contrast, there was a notable decline in CD34^-^CD7^-^CD4^+^ cell counts in HIV^+^ samples compared to those in HIV^-^ samples at week 3 (Fig. 2C). Further analysis of CD34^-^ cells revealed that the CD7^+^CD4^+^ cells contained CD4/CD8 double-positive cells and CD3^+^CD4^+^ cells, whereas the CD7^-^CD4^+^ cells were mostly CD3^+^CD4^+^ cells (Extended Data Fig. 3). In the analysis of CD34^-^CXCR4^+^, CD34^-^CD7^+^CXCR4^+^ and CD34^-^CD7^-^CXCR4^+^ cells, the CD34^-^CD7^-^CXCR4^+^ cells exhibited the greatest growth impairment following HIV-1 infection (Extended Data Fig. 4).

**Figure 2.**
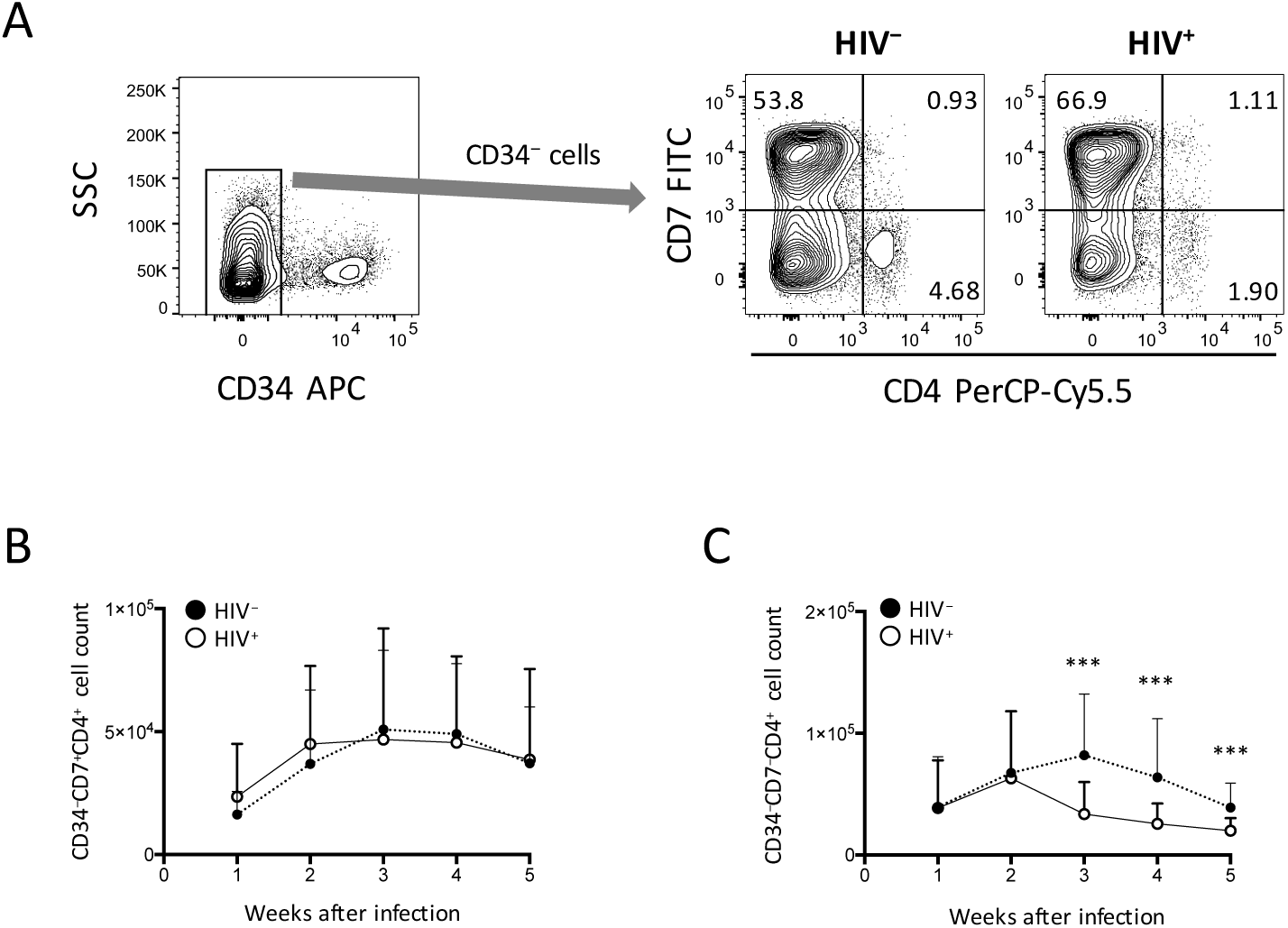
Pre-exposure of primary cord-derived CD34^+^ cells to HIV-1 affected the dynamics of OP9-DL1 cocultured cells (n = 12). (A) Representative plots showing the loss of CD34^-^CD7^-^CD4^+^ cells in the HIV-infected cocultures 3–5 weeks after infection. (B-C) Cell counts were compared between HIV^+^ and HIV^-^ cocultures for (B) CD34^-^CD7^+^CD4^+^ and (C) CD34^-^CD7^-^CD4^+^ cells. Statistical analyses were performed using the Wilcoxon matched-pairs signed rank test. ***: *P* < 0.001.

**Figure 3.**
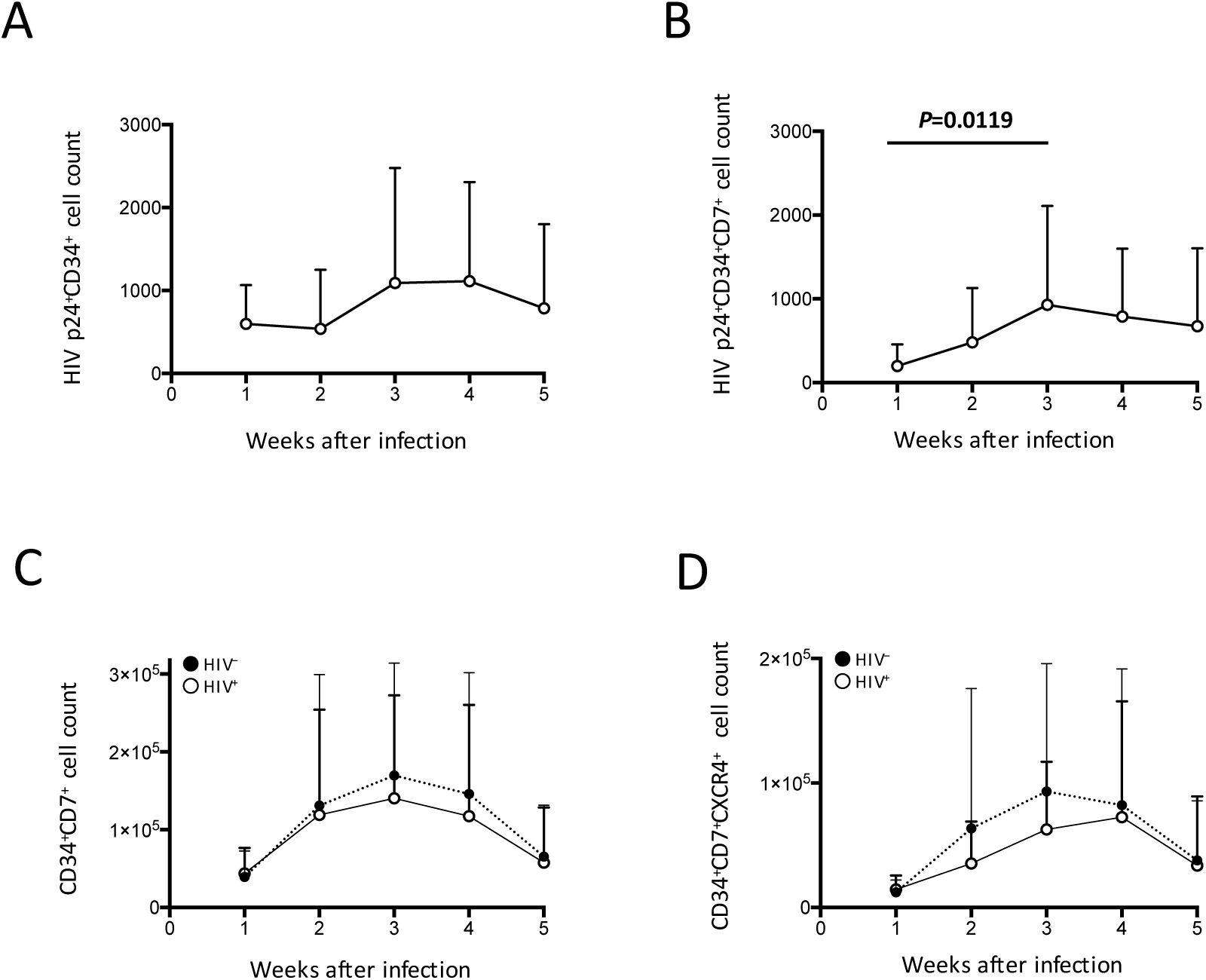
Pre-exposure of primary cord-derived CD34^+^ cells to HIV-1 affected the dynamics of CD34^+^ cells in the subsequent OP9-DL1 cocultures (n = 12). (A) HIV p24^+^CD34^+^ cell counts in HIV^+^ OP9-DL1 cocultures. (B) HIV p24^+^CD34^+^CD7^+^ cell counts in HIV^+^ OP9-DL1 cocultures. (C–D) Cell counts were compared between HIV^+^ and HIV^-^ cocultures for (C) CD34^+^CD7^+^ and (D) CD34^+^CD7^+^CXCR4^+^ cells. Statistical analyses were performed using the Wilcoxon matched-pairs signed rank test.

### The dynamics of CD34^+^ cells in OP9-DL1 cocultures with HIV–pre-exposed CD34^+^ cells

HIV infection of CD34^+^ cells was detected by intracellular p24^+^ staining followed by flow cytometric analysis (Fig. 3A–B, Extended Data Fig. 5A–D). p24^+^CD34^+^ cells failed to grow by week 2 (Fig. 3A), in contrast to the rapid growth of p24^+^ cells at this time point (Fig. 1C). p24^+^CD34^+^CD7^+^ cell counts increased at weeks 2 and 3 (Fig. 3B) in association with a low average CD34^+^CD7^+^ cell count in week 1 after infection and a rapid increase in week 2 (Fig. 3C). Then, CD34^+^ cell dynamics in HIV^+^ samples were compared with those in HIV^-^ samples (Fig. 3C-D, Extended Data Fig. 5E–G). There were non significant differences in CD34^+^CD7^+^ cell counts between HIV^+^ and HIV^-^ cocultures (Fig. 3C). The difference in the average frequencies of CD34^+^CD7^+^ cells was greatest at week 3 (Fig. 3C). However, the average CD34^+^CD7^+^CXCR4^+^ cell count in HIV^+^ cocultures versus that in HIV^-^ cocultures exhibited the greatest difference at week 2 (Fig. 3D). Therefore, CD34^+^CD7^+^CXCR4^+^ cells may be affected by HIV-1 earlier than the entire CD34^+^CD7^+^ cell population or cells of other phenotypes.

### HIV-1_NL4-3_ infection of HSPC-derived cells after 4–6 weeks of coculture with OP9-DL1 cells

To better describe the effect of HIV infection on the short-term dynamics of CD34^+^ cells, experiment 2 was performed (short coculture, n = 12, Fig. 4A). The experiment 2 results are shown in Figs. 4–6, Extended Data Figs. 6–8 and Extended Data Tables 1–2. Briefly, coculture of OP9-DL1 cells with HIV-uninfected cord-derived CD34^+^ cells for 4–6 weeks produced a mixture of cells with different phenotypes as determined by the expression of CD34, CD4, CD8, CD7 and CXCR4 (Figs. 1–3, Extended Data Figs. 1–5). CD4 expression levels in CD34^+^ cells before infection were lower than those in CD34^-^ cells (Fig. 4B). Cells were then harvested and infected with HIV-1_NL4-3_ following the procedures for HIV infection of CD34^+^ cells (Fig. 1A). The infected cells were cocultured again with a new OP9-DL1 monolayer and further incubated for 1 week (Fig. 4A). One week after infection, HIV replication was detected in all HIV-infected samples (Fig. 4C). The majority of HIV-1 p24^+^ cells were CD34^-^(Fig. 4C, Extended Data Fig. 6). The majority of p24^+^CD34^+^ cells were CD7^+^ (Fig. 4C, Extended Data Table 1).

**Figure 4.**
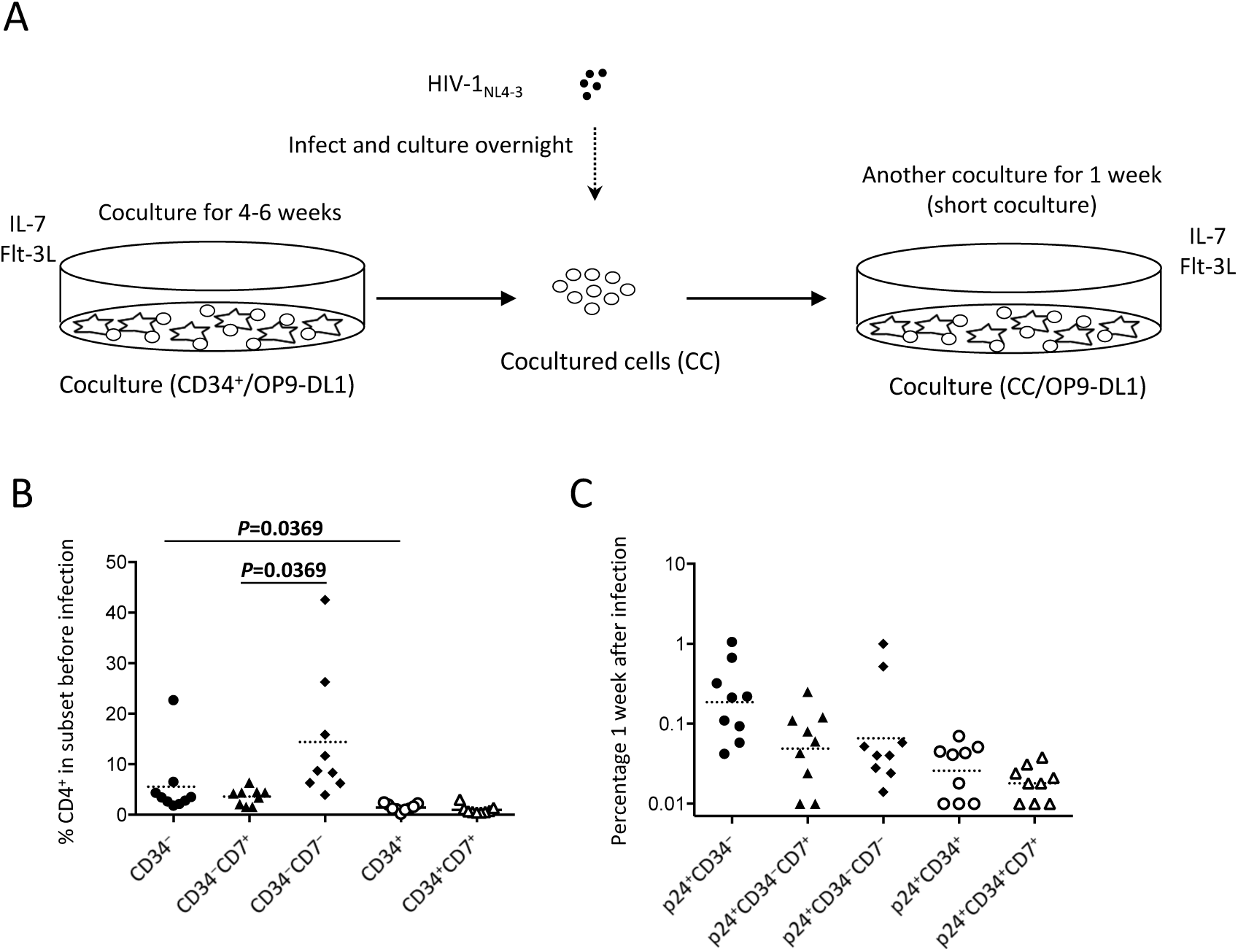
Experiment 2 (short coculture, n = 9) was performed to further analyse the dynamics of CD34^+^ cells in the presence of HIV-1. (A) Schematic representation of experiment 2 (short coculture). Primary cord-derived CD34^+^ cells were cocultured with OP9-DL1 cells for 4–6 weeks. Cells were then collected and infected with HIV-1NL4-3. The infected cells were cocultured again with OP9-DL1 cells for 1 week, collected and analysed. (B) Percentages of CD4^+^ cells in CD34^-^, CD34^-^CD7^+^, CD34^-^CD7^-^, CD34^+^ and CD34^+^CD7^+^ subsets before infection. A multiple comparison test was performed to compare CD34^-^ and CD34^+^ cells or CD34^-^CD7^+^ and CD34^-^CD7^-^ cells. (C) Frequencies of HIV p24^+^, p24^+^CD34^-^CD7^+^, p24^+^CD34^-^CD7^-^, p24^+^CD34^+^ and p24^+^CD34^+^CD7^+^ cells measured 1 week after the second post-infection coculture (n = 9). Statistical analysis was performed using the multiple comparison test with Dunn’s method.

### Partial loss of CD34^+^CD7^+^CXCR4^+^ cells after HIV-1_NL4-3_ infection of OP9-DL1– cocultured cells

The phenotypes of the cells in experiment 2 (short coculture) were analysed 1 week after infection (Fig. 5, Extended Data Fig. 7, Extended Data Table 2). The whole cell counts were obtained using parts of the tested batches, with no significant differences noted between HIV^+^ and HIV^-^ samples (n = 4, *p* = 0.5000, Extended Data Table 2). The frequencies of CD4^+^CD8^+^, CD34^+^ and CD34^+^CD7^+^ cells were not significantly different between HIV^+^ and HIV^-^ cocultures (Extended Data Fig. 7). Regarding CXCR4 expression levels in different subsets, the frequencies of CD34^+^CD7^+^CXCR4^+^ cells in HIV^+^ samples were significantly reduced compared to those in HIV^-^ samples (Fig. 5A and C middle). This may be in accordance with the results in Fig. 3D in which the growth of CD34^+^CD7^+^CXCR4^+^ cells slowed earlier (at week 2) than those of other phenotypes (at week 3).

**Figure 5.**
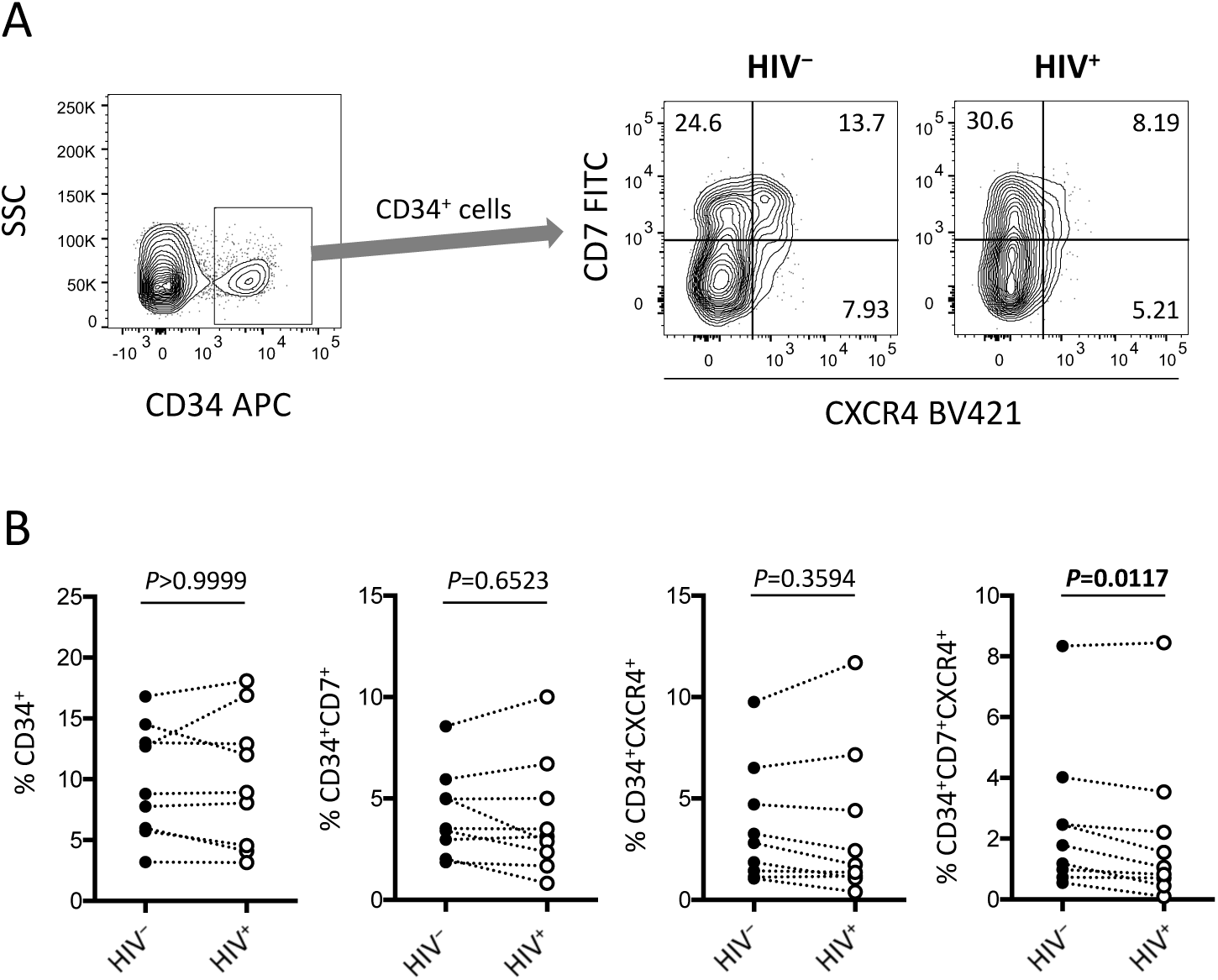
CD34^+^ cells in experiment 2 (short coculture, n = 9) were analysed 1 week after the second post-infection coculture. (A) Representative plots showing CD7/CXCR4 expression levels in CD34^+^ cells. An HIV^+^ sample is compared to its autologous uninfected counterpart. (B) Frequencies of CD34^+^ (left), CD34^+^CD7^+^ (middle left), CD34^+^CXCR4^+^ (middle right) and CD34^+^CD7^+^CXCR4^+^ (right) cells were compared between HIV^+^ and HIV^-^ samples. Statistical analyses were performed using the Wilcoxon matched-pairs signed rank test.

### Increased number of CD34^+^CD7^+^CD4^+^ cells after HIV-1_NL4-3_ infection of OP9-DL1– cocultured cells

Figure 5 indicates the distinct dynamics of CD34^+^CD7^+^CXCR4^+^ cells in the presence of HIV-1. For increased clarity, CD34^+^CD7^+^ cells were further analysed for CD4 expression (Fig. 6, Extended Data Fig. 8). Surprisingly, HIV-infected samples had higher frequencies of CD34^+^CD7^+^CD4^+^CXCR4^+^ cells than uninfected samples (Fig. 6A and B). These cells were found to be partially HIV p24^+^ after carefully adjusting the compensation matrices of the flow cytometry data (Fig. 6A and C). One of nine batches was selected for further analyses of CD34^+^CD7^+^CD4^+^ cells. Both the frequencies and CD4 fluorescence intensities of CD34^+^CD7^+^CD4^+^ cells were higher in HIV^+^ samples than in HIV^-^ samples (Extended Fig. 8B), implying the emergence of CD34^+^CD7^+^CD4hi cells. In addition, the frequencies of CD34^+^CD7^+^CD4^+^CXCR4^+^ cells were correlated with those of p24^+^ cells (Fig. 6D). A separate staining of an experiment 2 sample with annexin V at week 1 after infection didn’t result in increased annexin V reactivity of CD34^+^CD4^+^ cells in HIV^+^ samples compared to those in HIV^-^ samples (data not shown).

**Figure 6.**
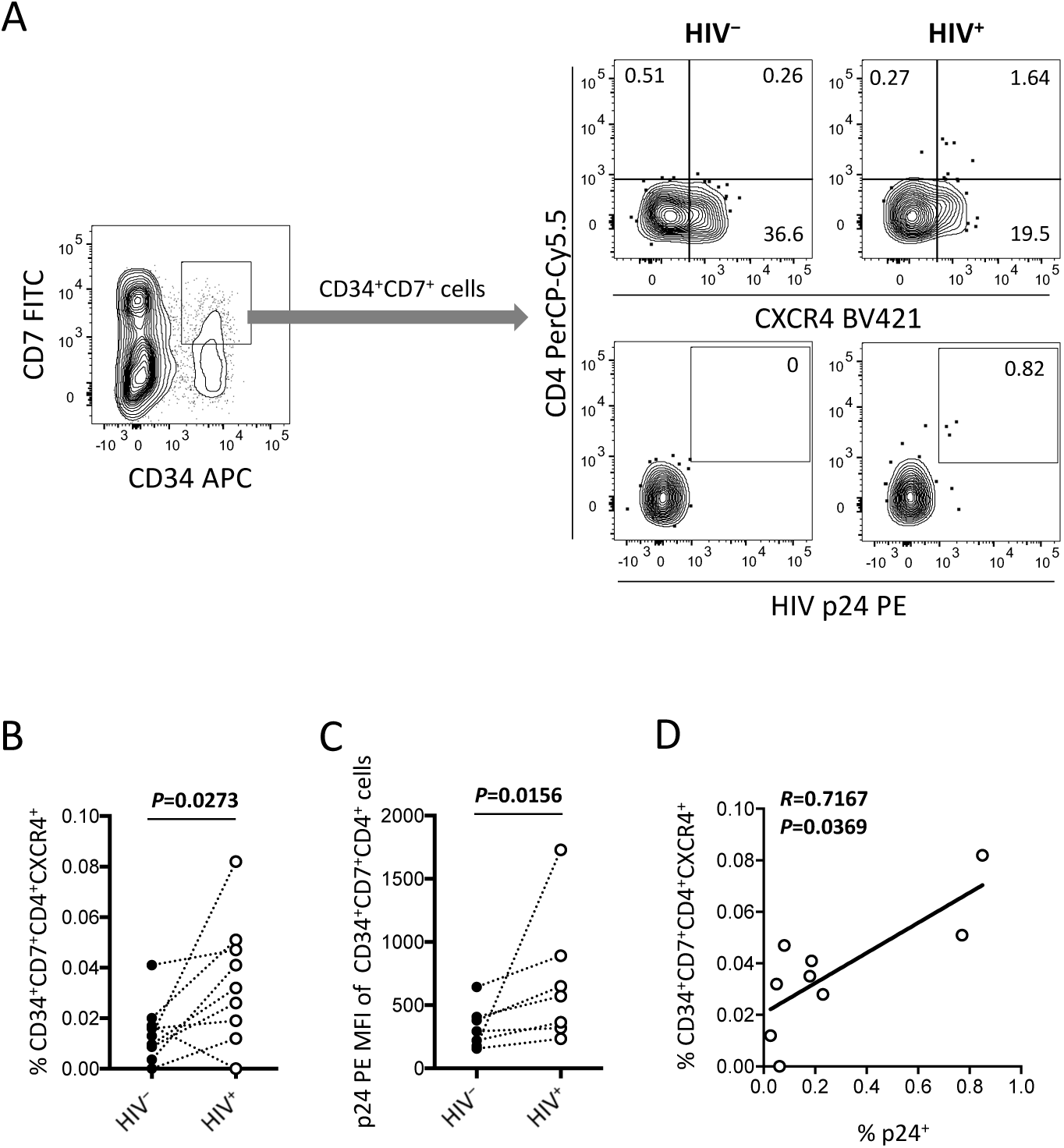
CD34^+^CD7^+^ cells in experiment 2 (short coculture, n = 9) were analysed 1 week after the second post-infection coculture. For these analyses, compensation matrices were checked carefully to exclude false-positive events. (A) Representative plots showing CD4/CXCR4 and CD4/p24 expression levels in CD34^+^CD7^+^ cells. An HIV^+^ sample is compared to its autologous uninfected counterpart. (B) Comparison of the frequencies of CD34^+^CD7^+^CD4^+^CXCR4^+^ cells between HIV^+^ and HIV^-^ samples. (C) p24 mean fluorescence intensities (MFIs) of CD34^+^CD7^+^CD4^+^ cells were compared between HIV^+^ and HIV^-^ samples (n = 7). (D) The frequencies of CD34^+^CD7^+^CD4^+^CXCR4^+^ cells were correlated with those of p24^+^ cells. Comparisons were performed using the Wilcoxon matched-pairs signed rank test. Spearman’s correlation coefficient was calculated for the correlation analysis.

## Discussion

Regarding the dynamics of CD34^+^ cells in HIV-infected individuals, there has been a debate regarding whether HSPCs comprise an unignorable viral reservoir ^26^. There is also a possibility of modified turnover of HSPCs through HIV infection and depletion of CD4^+^ cells. For example, HIV infection commonly manifests as bone marrow abnormalities ^7^. Some patients fail to exhibit CD4^+^ T-cell recovery even after effective antiretroviral therapy, thus being termed immunological nonresponders. Such immunological nonresponsiveness can be associated with immune activation and/or bone marrow impairment ^27,28^. To approach these problems, the present study revealed a method for utilising the OP9-DL1 coculture system, which enables in vitro follow-up of the early events in T-cell differentiation such as lymphoid progenitor cell generation that normally occur in the bone marrow as well as CD4^+^ thymocyte differentiation in the thymus. The events observed in the cocultures of OP9-DL1 and human CD34^+^ cells are likely to involve interactions between SDF-1 and CXCR4, as mouse SDF-1 expressed by OP9-DL1 cells has high amino acid sequence identity with human SDF-1 ^29^. The observation is in accordance with previous reports illustrating that the SDF-1/CXCR4 pair is crucially involved in the homing and repopulation of HSPCs in specific bone marrow niches ^30^ and the entire T-cell developmental process in the thymus ^21,31^. It has been reported that HSPCs and thymocytes express CXCR4, but their CCR5 expression is limited ^32^. Another report indicated that HIV utilises CXCR4 when it infects multipotent progenitor cells ^33^. CXCR4-tropic HIV strains might contribute to pathogenesis by interfering with haematopoiesis and/or lymphopoiesis ^34-36^, although the underlying mechanisms are yet to be unveiled.

The present results provide insights into the impact of HIV-1 on T-lineage differentiation of haematopoietic progenitor cells. To be more physiological than the previous studies analysing the influence of HIV on CD34^+^ cells, the present study didn’t include strong stimuli such as 50–100 ng/mL of stem cell factor, thrombopoietin, FMS-like tyrosine kinase 3 ligand or interleukin 3 ^33,37^. In experiment 1 (long coculture), HIV-infected samples exhibited similar CD4^+^ cell production rates as uninfected samples at weeks 1 and 2 but reduced CD4^+^ cell production rates from week 3 to week 5 (Fig. 2). This was most clearly observed with the dynamics of CD34^-^CD7^-^CD4^+^ cells (Fig. 2C) that were mostly CD3^+^ T cells (Extended Data Fig. 3). It is unclear how the reduction of cell growth at week 3 after infection was so accurately reproduced in the 12 samples tested (Fig. 2C, individual data not shown). However, in the subsequent analyses of CD34^+^ cells, the growth rates of CD34^+^CD7^+^CXCR4^+^ cells tended to fall as early as week 2 (Fig. 3D). This was observed in 7/12 samples tested (data not shown), and the decline occurred earlier than the reduction in CD34^-^CD7^-^CD4^+^ cell counts (week 3). Because CD34^+^CD7^+^ cells represent lymphoid progenitor cells ^38,39^, their involvement in defining the production rates of CD4^+^ cells may be speculated.

Subsequently, experiment 2 (short coculture, Figs. 4–6) was conducted to examine the effects of HIV-1 on the short-term dynamics of T-lineage cells including CD34^+^ progenitors. HIV-1 infection resulted in significant decreases of the frequency of CD34^+^CD7^+^CXCR4^+^ cells 1 week after infection (Fig. 5B right), which may be consistent with the results of experiment 1 (Fig. 3D). Because of the limitation of the present study, the fate of the lost CD34^+^CD7^+^CXCR4^+^ cells in the HIV^+^ cocultures is unknown. The differentiation capacity of the remaining CD34^+^CD7^+^CXCR4^+^ cells also remains untested. Interestingly, a recent article reported that bone marrow CD34^+^progenitor cells from HIV-infected patients exhibit impaired T-cell differentiation potential ^40^, which was related to proinflammatory cytokines. Cytokine analyses may be applicable to the coculture assays established in the present study.

The CD34^+^CD7^+^ progenitors were further analysed to better understand their association with HIV-1 infection. Interestingly, the frequencies of CD34^+^CD7^+^CD4^+^ cells were elevated in HIV-infected samples compared to those in uninfected samples (Fig. 6, Extended Data Fig. 8). In addition, CD34^+^CD7^+^CD4^+^ cells were found to be partially HIV p24^+^ (Fig. 6A and C). It may be fascinating to interpret the results as CD4 upregulation following HIV infection, thereby driving the differentiation of T-lineage precursor cells. However, caution may be necessary because exclusion of possible CD34 upregulation in the infected CD4^+^ cells was not confirmed in this study (Extended Data Fig. 8), although the flow cytometry data did not indicate correlations between HIV p24 expression and CD34 upregulation in CD34low cells (data not shown). Further studies may be designed to test shorter coculture periods than 1 week and/or investigate the association of HIV-1 infection with factors that regulate the expression of CD3, CD4, CD7 and CD34. Such analyses will also help better clarify the debate on the issues of CD34^+^ viral reservoirs ^26^.

The findings in this article highlight the potential of anti-HIV treatments such as gene therapy using CD34^+^ HSPCs followed by transplantation because in this manner, all haematopoietic events in the host can be protected against HIV infection by the gene products even in the absence of effective immune responses ^41^. The problem is that although CCR5 may be considered a reasonable target for knockout or knockdown to prevent infection by CCR5-tropic HIV strains, CXCR4 expression should not be altered in HSPCs because of the essential biological functions of the molecule ^42^. Therefore, instead of modulating CXCR4 expression, anti-HIV modalities targeting an HIV gene or component may be desirable for protecting haematopoietic cells including T-lineage cells from CXCR4-tropic HIV infection ^43^.

In summary, the results of the long/short coculture of human CD34^+^ cells and derivatives with OP9-DL1 cells in the presence of HIV-1 indicate that the dynamics of CD34^+^CD7^+^ lymphoid progenitors may be affected more quickly by HIV-1 than CD34^-^CD4^+^ thymocytes and T cells despite the lower susceptibility of CD34^+^CD7^+^ cells to HIV-1 infection as suggested by their lower CD4 expression levels. Further studies may be designed to elucidate the underling mechanisms. In addition, the preceding reduction of CD34^+^CD7^+^CXCR4^+^ cell growth at week 2 followed by the reduction of CD34^-^CD4^+^ cell growth at week 3 in the HIV^+^ cocultures may illustrate the mechanism by which reduced sizes of progenitor cell pools may decelerate the production of T cells in HIV-infected patients (Extended Data Fig. 9). This may occur in combination with different mechanisms of CD4^+^ T-cell depletion including direct cytopathic effects, apoptosis and antigen-specific immunological mechanisms ^44-46^.

## Methods

### Virus stocks

Stocks of HIV-1_NL4-3_ were produced via transfection of 293T cells with the molecular clone DNA p_NL4-3_ ^47^. After transfection, the culture supernatant was collected, and viral loads were determined using an HIV p24 enzyme-linked immunosorbent assay (ELISA) kit (ZeptoMetrix, NY, USA).

### Cells

Umbilical cord blood samples were collected at Fukuda Hospital, Kumamoto, Japan after obtaining informed consent. Cord blood mononuclear cells were isolated using Pancoll (PAN-Biotech GmbH, Aidenbach, Germany) and centrifugation at 800 rpm for 20 min. CD34^+^ cells were labelled and selected using human CD34 microbeads and LS columns (Miltenyi Biotec, NSW, Australia). The purity constantly exceeded 92%. The OP9-DL1 cell line was generated via retroviral transduction of the OP9 cell line (RCB1124, Riken, Tsukuba, Japan) with human DL1. The cell line was maintained in α-MEM medium (Wako Pure Chemical Industries, Osaka, Japan) supplemented with 10% heat inactivated fetal bovine serum (FBS, GE Healthcare, Tokyo, Japan).

### Antibodies

Anti-human CD8 Brilliant Violet (BV) 510 (clone RPA-T8), anti-human CD3 PE-Cy7 (clone UCHT1) and anti-human CD34 APC (clone 8G12) were purchased from BD Biosciences (NSW, Australia). Anti-human CD4 PE-Cy7 (clone OKT4), anti-human CD4 PerCP-Cy5.5 (clone OKT4) and anti-human CXCR4 BV421 (clone 12G5) were purchased from BioLegend (CA, USA). Anti-human CD3 ECD (clone UCHT1) and anti-HIV-1 p24 PE (clone FH190-1-1, also known as KC57 RD1) were purchased from Beckman Coulter (NSW, Australia). Anti-human CD7 FITC (clone CD7^-^6B7) was purchased from CALTAG Laboratories (CA, USA).

### Coculture of human cells with OP9-DL1

The OP9-DL1 coculture experiment was performed following a previously published protocol with modifications ^48^. Briefly, 2 × 105 HIV-infected or uninfected cord-derived CD34^+^ cells were seeded in a 6-well plate (Corning, VIC, Australia) containing a OP9-DL1 monolayer. The coculture was maintained for 5 weeks in α-MEM medium supplemented with 20% heat inactivated FBS, 5 ng/mL recombinant human FMS-like tyrosine kinase 3 ligand (R&D Systems, MN, USA) and 5 ng/mL recombinant human interleukin 7 (Miltenyi Biotec). Cells were passaged weekly with vigorous pipetting and filtering through a 70-µm membrane and cocultured again with a fresh monolayer of OP9-DL1 cells. A portion of the collected cells was analysed by flow cytometry.

## HIV infection

The method for in vitro HIV infection of CD34^+^ cells was described previously ^49^. To infect primary cord-derived CD34^+^ cells with HIV-1NL4-3, a 48-well plate (Corning) was treated overnight with RetroNectin (Takara Bio, Kusatsu, Japan) at a concentration of 10 µg/mL. CD34^+^ cells were re-suspended in the OP9-DL1 medium, seeded at 2 × 105 per well in the coated plate and infected with 200 ng (p24) of HIV-1_NL4-3_ using spinoculation at 1200 × *g* at 34°C for 30 min. Cells were further cultured overnight and cocultured in a 6-well plate with a fresh monolayer of OP9-DL1 cells. For HIV infection of OP9-cocultured human cells, the cell concentration was modified to 5 × 10^5^ per well.

### Flow cytometry

Surface and intracellular antigen expression was analysed using a FACS LSR II (BD Biosciences), FACS Diva v6.0 software (BD Biosciences) and FlowJo v10.3 software (FlowJo, OR, USA). Dead cells were discriminated using Live/Dead Fixable Near-IR Dead Cell Stains (Thermo Fisher Scientific, VIC, Australia). Live cells were further gated to exclude doublets from the analysis by plotting FSC-H and FSC-A.

### PCR analysis of HIV DNA

Cellular DNA was extracted using a Kaneka Easy DNA Extraction Kit (Kaneka, Takasago, Japan). DNA extraction was followed by the PCR analysis using an HIV *gag* primer set (sense: 5'-AGTGGGGGGACATCAAGCAGCCATGCAAAT-3', antisense: 5'-TACTAGTAGTTCCTGCTATGTCACTTCC-3') as described previously ^50^. The oligo DNAs were purchased from Sigma-Aldrich (Tokyo, Japan).

### Statistical analysis

Statistical analysis was performed using GraphPad Prism software version 7.0 (GraphPad Software, CA, USA). Statistical significance was defined as *P* < 0.05. Comparisons between HIV-infected and uninfected samples were performed using Wilcoxon’s matched-pairs signed rank test, unless otherwise stated. Multiple comparison analyses were done, if necessary, using Dunn’s method. Spearman’s correlation coefficients were calculated for correlation analyses.

### Data availability

The data that support the findings of this study are available from the corresponding author upon reasonable request.

## Acknowledgements

This work was supported by the National Health and Medical Research Council, Australia (project grant). It was also partially supported by grants from the following organizations: the Japan Agency for Medical Research and Development (Research program on HIV/AIDS, No. 16fk0410108h0001); Ministry of Education, Culture, Sports, Science and Technology, Japan (Grants-in-Aid for Science Research, No. 25114711) and Kumamoto University (AIDS International Collaborative Research Grant). I thank Drs. Kazuo Matsui and Shoichi Kawakami of Fukuda Hospital, Kumamoto, Japan for their assistance with cord blood sampling. I thank Prof. Seiji Okada of Center for AIDS Research, Kumamoto University and his staff for their assistance with preparing resources. I thank Prof. Anthony D Kelleher and Dr. Kazuo Suzuki of the Kirby Institute for infection and immunity in society, UNSW Australia for their support. I thank Enago (www.enago.jp) for the English language review.

## Author information

The author declares no competing financial interests associated with the present work. Correspondence and requests for materials should be addressed to.

## Extended Data figures and tables

**Extended Data Figure 1.**
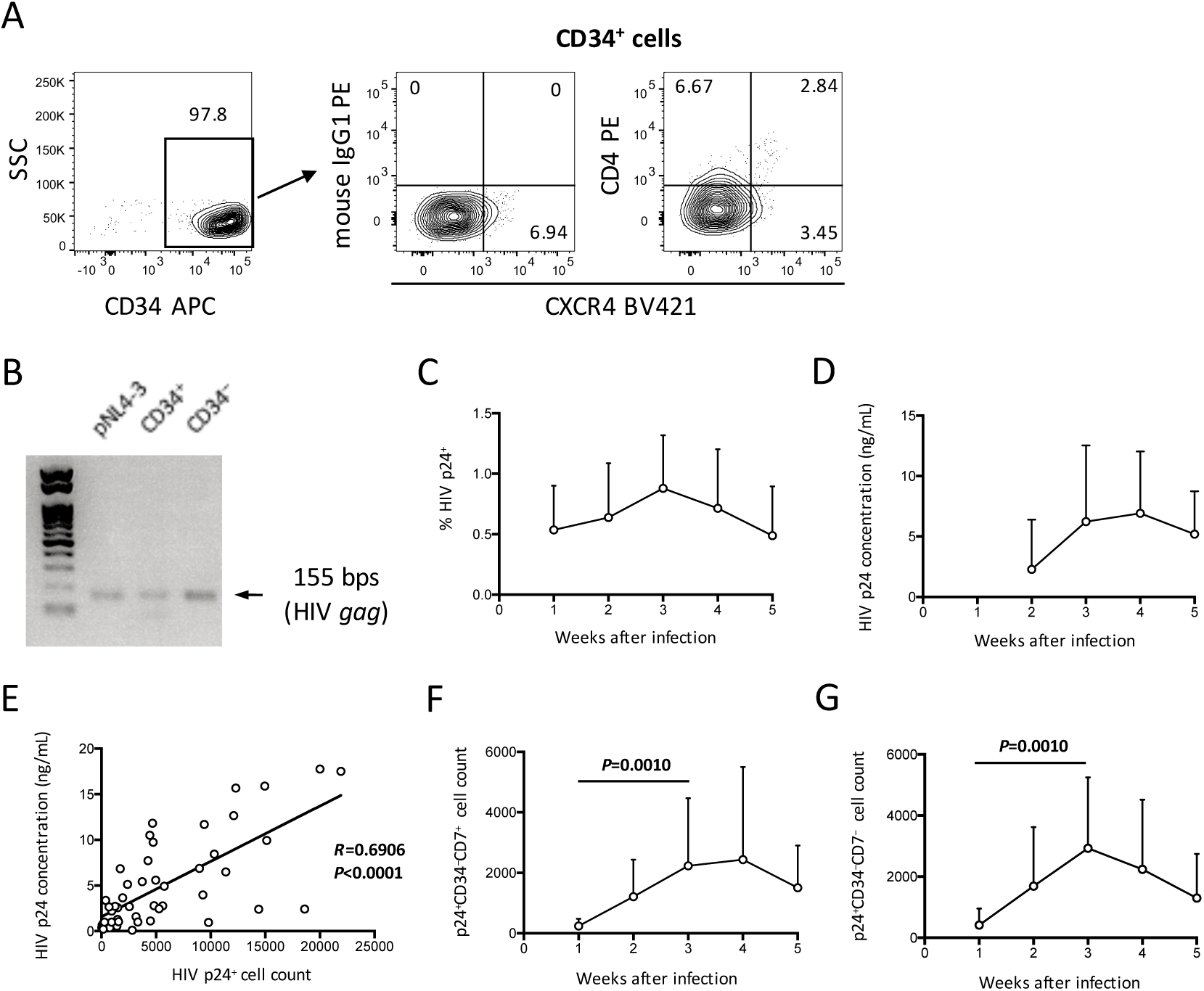
HIV infection of cord-derived CD34^+^ cells and derivatives were evaluated. (A) Primary cord-derived CD34^+^ cells were gated and tested for CD4 and CXCR4 expression by flow cytometry. Representative plots are shown. The CD4 signal levels were compared to the signal levels of a PE-conjugated mouse isotype control antibody. (B) The CD34^+^ and CD34^-^fractions of an HIV-infected sample were separated using the CD34 microbead method. Total DNA was isolated and analysed by PCR. HIV gag DNA was detected in both the CD34^+^ and CD34^-^fractions. A PCR sample using the HIV molecular plasmid p_NL4-3_ was used as a control. (C) Percentage of HIV p24^+^ cells in HIV^+^ OP9-DL1 coculture samples tested weekly by intracellular staining and flow cytometry. (D) Coculture supernatants of HIV-infected samples were tested for HIV p24 concentrations by ELISA at weeks 2– 5. (E) Correlation analysis between the frequencies of HIV p24^+^ cells and HIV p24 concentrations. The weeks 2–5 data in F and G are included (n = 48). Spearman’s correlation coefficients are shown. (F) p24^+^CD34^-^CD7^+^ cell counts. (G) p24^+^CD34^-^CD7^-^cell counts. Comparisons were performed using the Wilcoxon matched-pairs signed rank test.

**Extended Data Figure 2.**
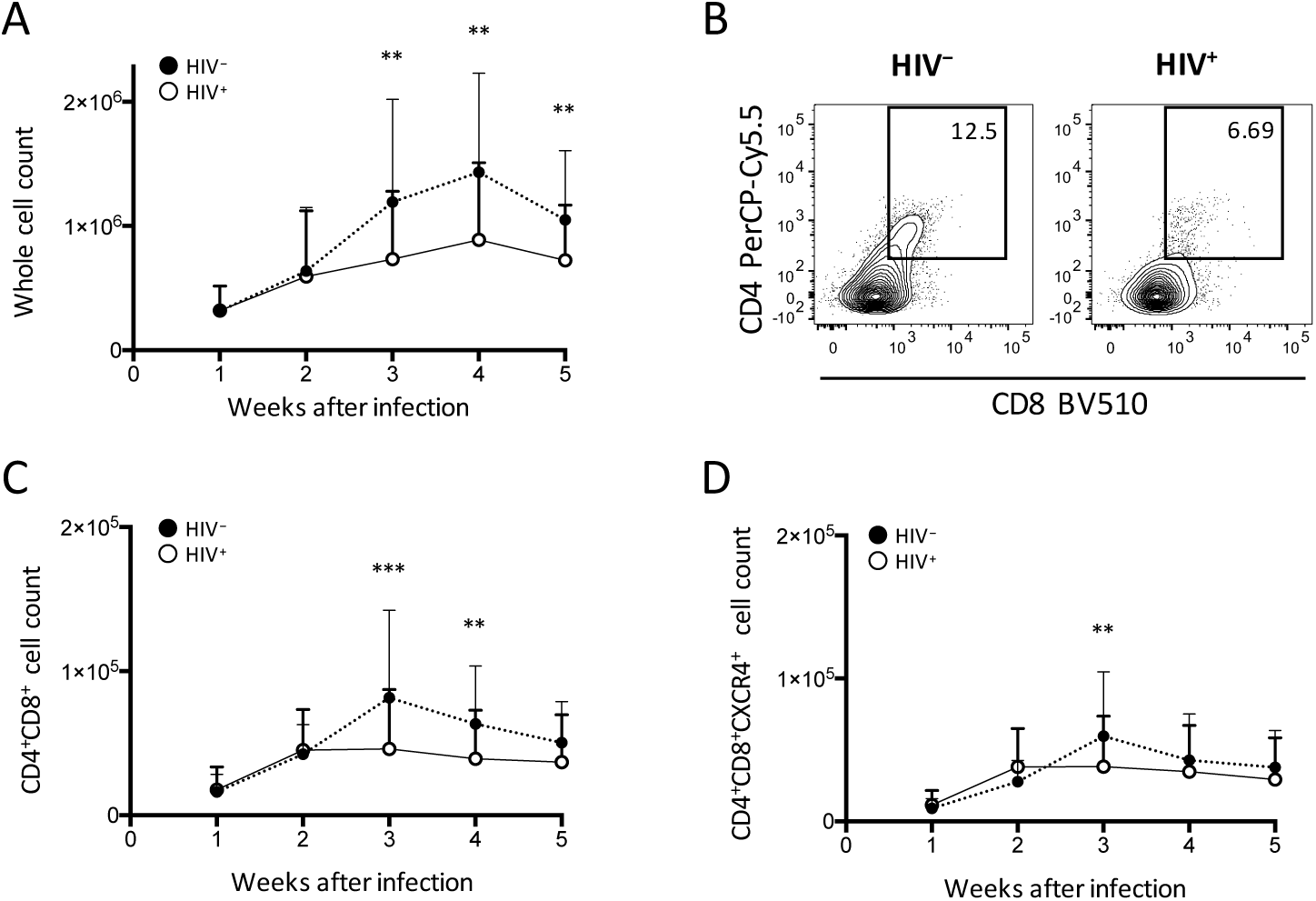
Pre-exposure of primary cord-derived CD34^+^ cells to HIV-1 affected the dynamics of OP9-DL1 cocultured cells. (A) Whole cell counts were compared between HIV^+^ and HIV^-^ cocultures. (B) Representative plots for samples displaying reduced frequencies of CD4^+^CD8^+^ cells 3–5 weeks after HIV pre-exposure of primary CD34^+^ cells and coculture. The plots were selected from week 4 samples. (C–D) Cell counts were compared between HIV^+^ and HIV^-^ cocultures for (C) CD4^+^CD8^+^ and (D) CD4^+^CD8^+^CXCR4^+^ cells. Statistical analyses were performed using the Wilcoxon matched-pairs signed rank test. **: *P* < 0.01, ***: *P* < 0.001.

**Extended Data Figure 3.**
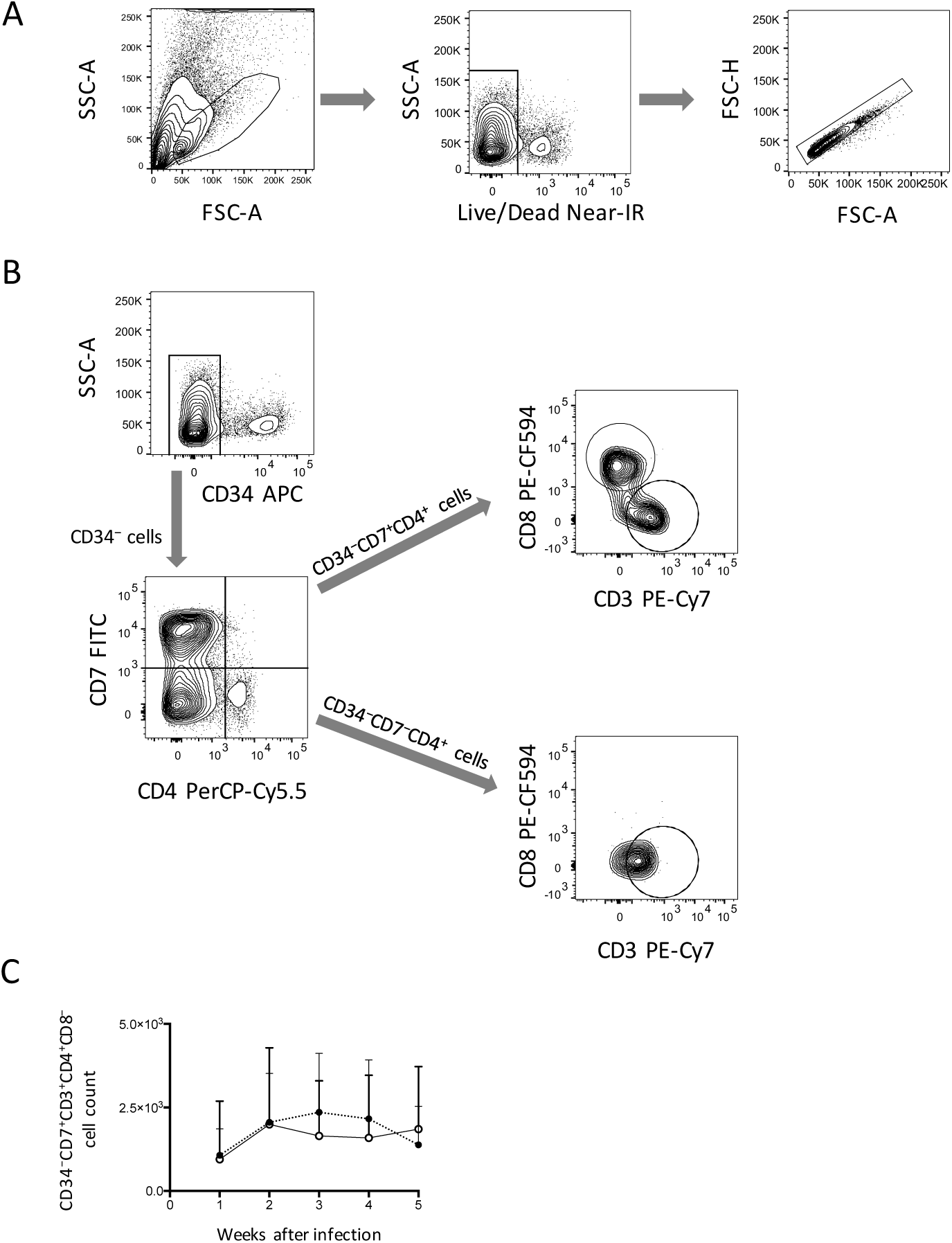
Gating strategies and analysis of CD34^-^CD4^+^ cells. (A) Flow cytometry plots showing the gates for lymphoid cells (left), live cells (middle), and singlet cells (right). (B) Flow cytometry plots showing the phenotype analysis of CD34^-^CD4^+^ cells in an OP9-DL1 coculture. Representative plots are shown. Cord-derived CD34^+^ cells were cocultured with OP9-DL1 cells for 4 weeks. Cells were collected and analysed by flow cytometry. CD34^-^ cells were gated by CD7 and CD4 expression. CD7^+^CD4^+^ and CD7^-^CD4^+^ cells were further gated by CD8 and CD3 expression. The analysis revealed that CD7^+^CD4^+^ cells contained different subsets including CD3-CD4^+^CD8^+^ and CD3^+^CD4^+^CD8-cells, whereas CD7^-^CD4^+^ cells were mostly CD3^+^CD4^+^CD8-cells. These results indicate that CD4/CD8 double-positive thymocytes, CD7^+^ CD4 T cells and CD7^-^CD4 T cells were generated via coculture of cord-derived CD34^+^ and OP9-DL1 cells. (C) CD34^-^CD7^+^CD3^+^CD4^+^CD8-cell counts.

**Extended Data Figure 4.**
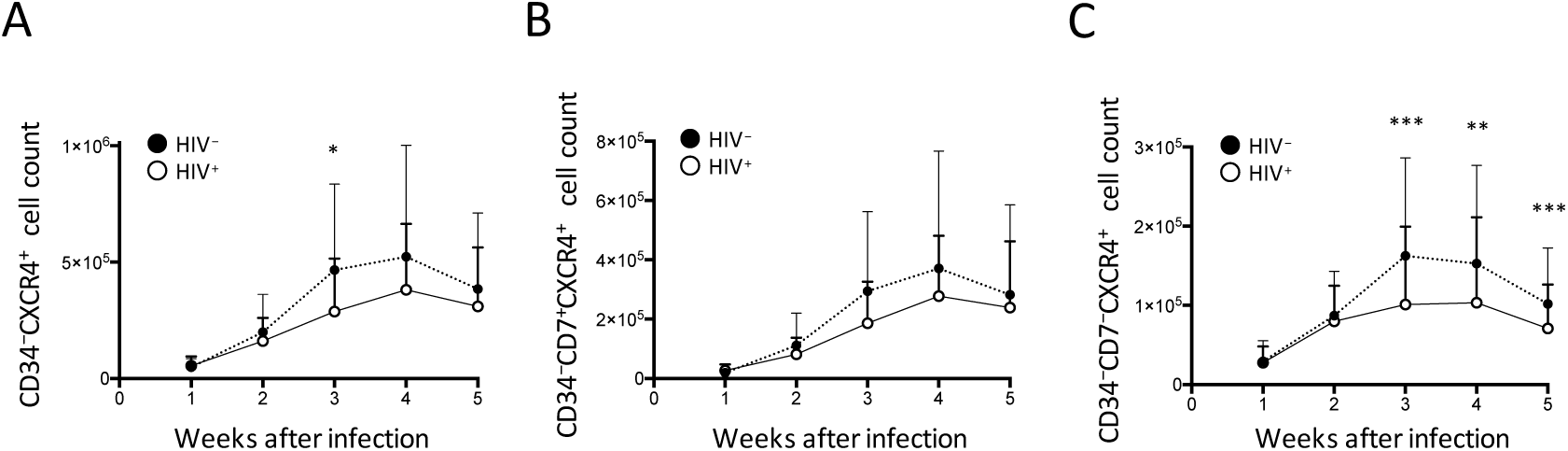
Phenotypes of CD34^-^ cells in the OP9-DL1 cocultures were analysed weekly for 5 weeks after HIV-1 infection. Cell counts were compared with those in uninfected samples. (A) CD34^-^CXCR4^+^ cell counts. (B) CD34^-^CD7^+^CXCR4^+^ cell counts. (C) CD34^-^CD7^-^CXCR4^+^ cell counts. Comparisons were performed using the Wilcoxon matched-pairs signed rank test. *: *P* < 0.05, **: *P* < 0.01, ***: *P* < 0.001.

**Extended Data Figure 5.**
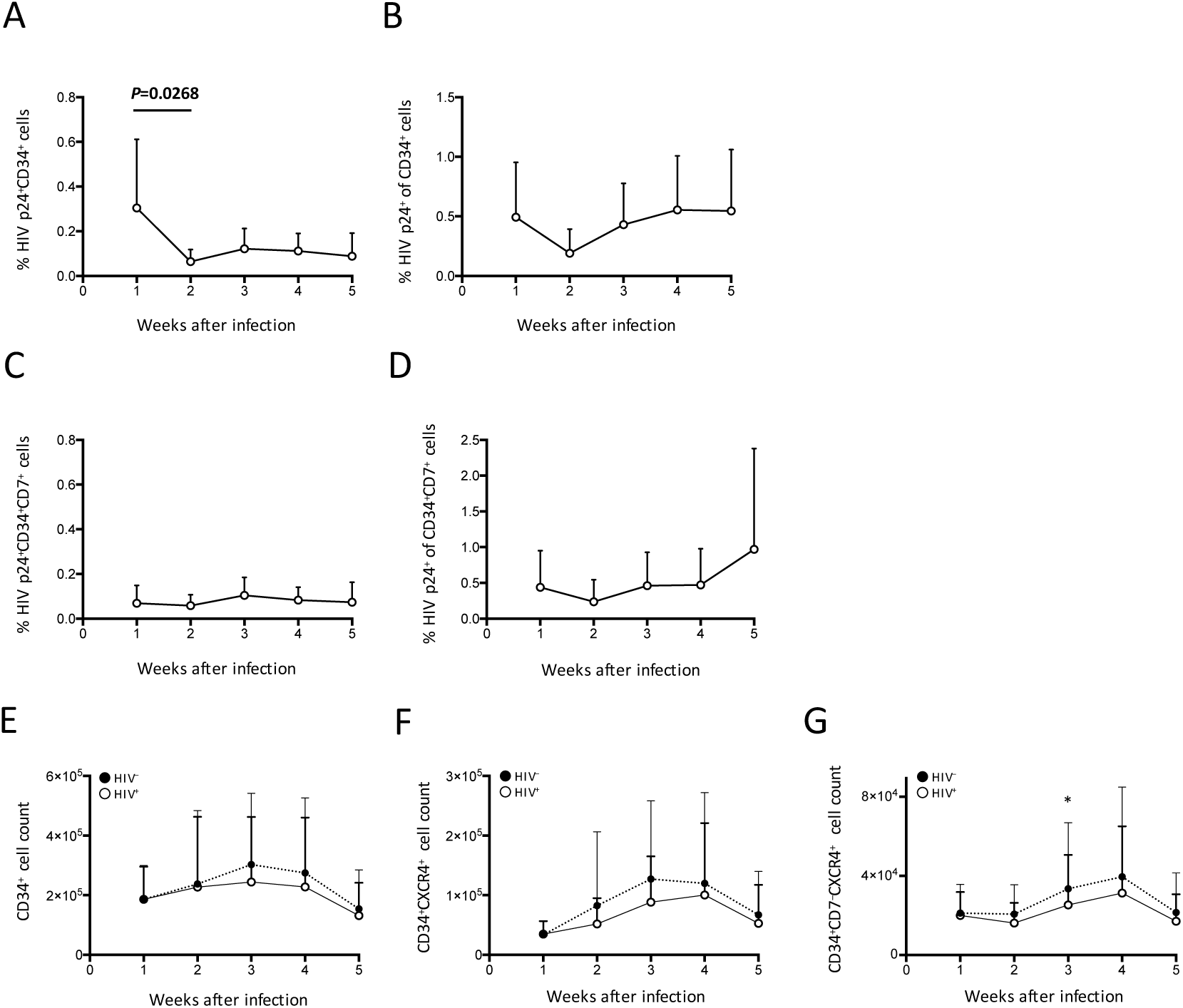
(A–D) HIV p24^+^ subset dynamics were analysed. (A) Percent p24^+^CD34^+^ cells. (B) Percent p24^+^ cells among CD34^+^ cells. (C) Percent p24^+^CD34^+^CD7^+^ cells. (D) Percent p24^+^ cells among CD34^+^CD7^+^ cells. (E–G) CD34^+^ subset dynamics were analysed and compared between HIV^+^ and HIV^-^ samples. (E) CD34^+^ cell counts. (F) CD34^+^CXCR4^+^ cell counts. (G) CD34^+^CD7^-^CXCR4^+^ cell counts.

**Extended Data Figure 6.**
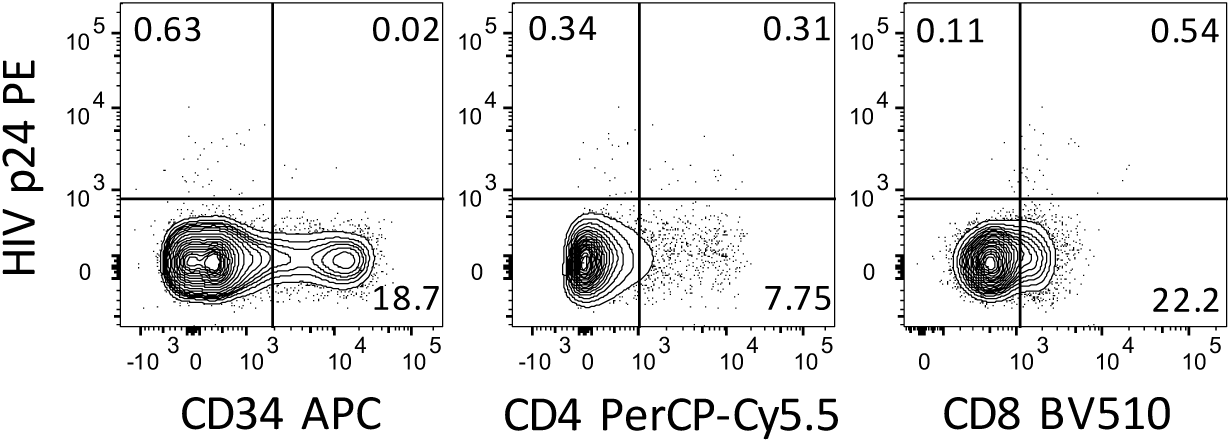
Experiment 2 (short coculture, n = 9) was performed to further analyse the dynamics of CD34^+^ cells in the presence of HIV-1. Representative plots showing HIV p24^+^ cells 1 week after infection. The majority of p24^+^ cells were CD34^-^(left). HIV replication may be causing CD4 downregulation (middle). Frequencies of CD8^+^ cells in p24^+^ cells were variable among the samples (right).

**Extended Data Figure 7.**
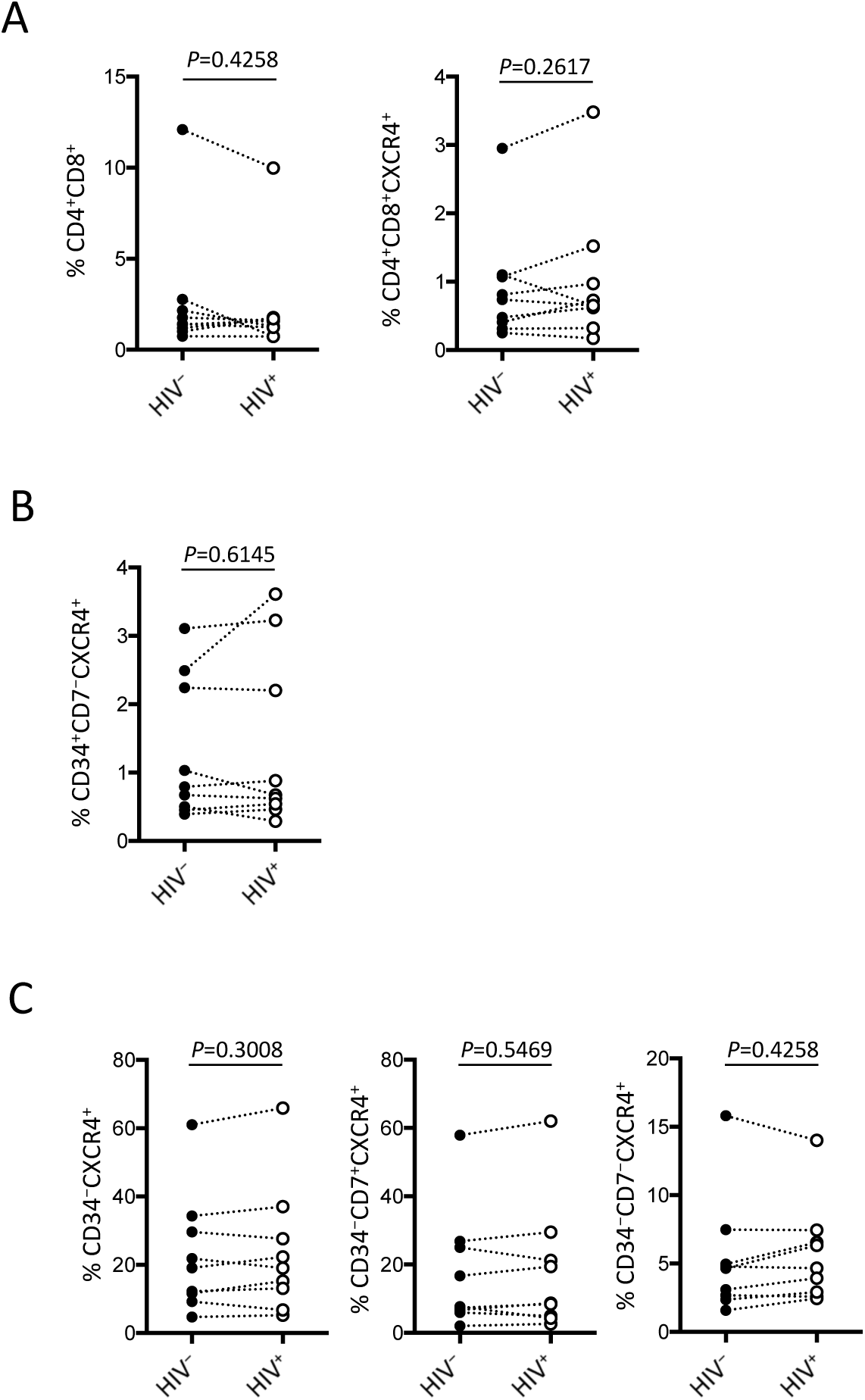
The frequencies of subsets were compared between HIV-infected and uninfected samples 1 week after infection. (A) CD4^+^CD8^+^ cells. (B) CD4^+^CD8^+^CXCR4^+^ cells. (C) CD34^-^CXCR4^+^ (left), CD34^-^CD7^+^CXCR4^+^ (middle) and CD34^-^CD7^-^CXCR4^+^ (right) cells. Comparisons were performed using the Wilcoxon matched-pairs signed rank test.

**Extended Data Figure 8.**
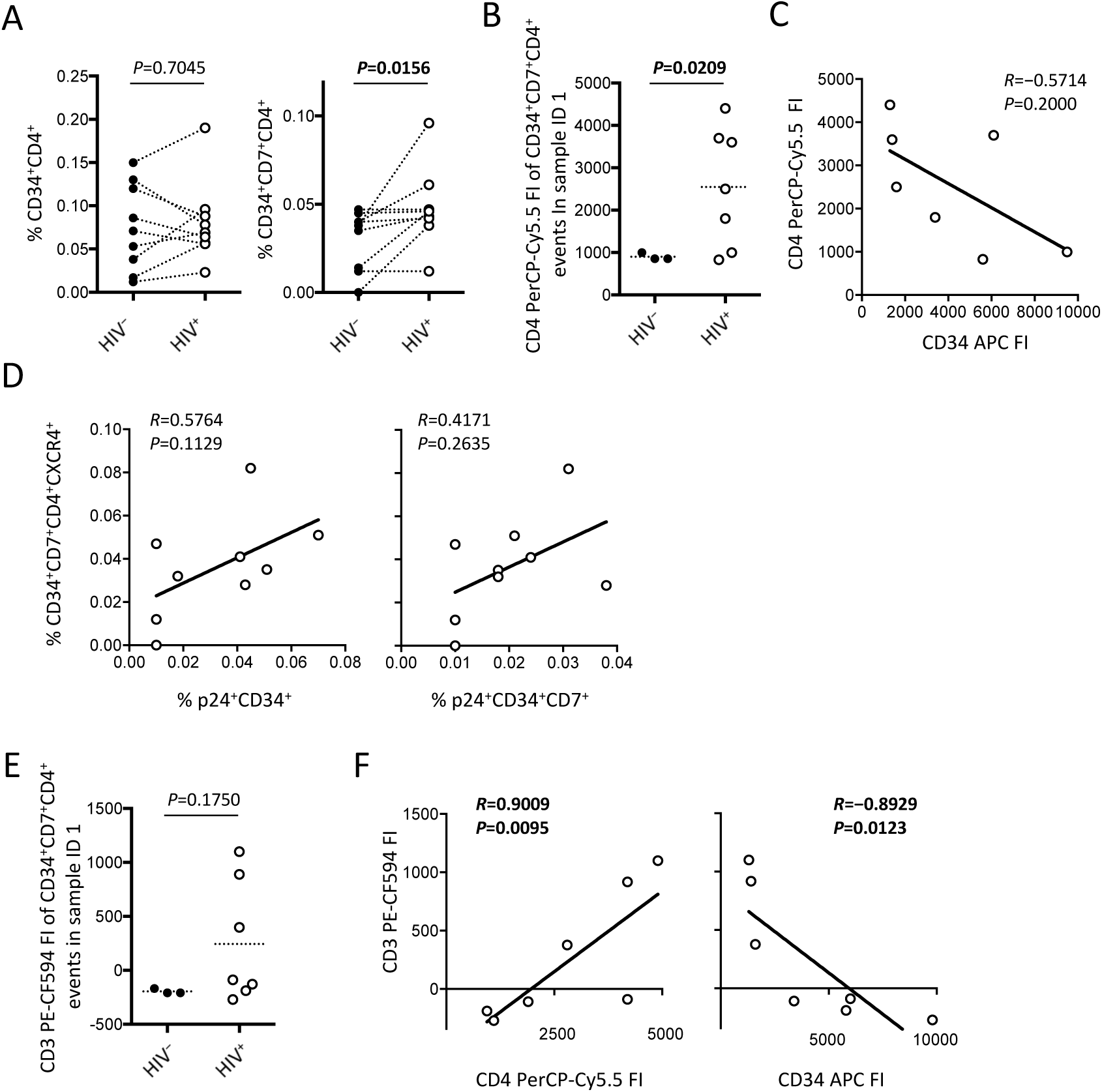
CD34^+^CD7^+^CD4^+^ cells were further analysed. (A) Comparison of CD34^+^CD4^+^ (left) and CD34^+^CD7^+^CD4^+^ (right) frequencies between HIV^+^ and HIV^-^ samples. (B) Fluorescence intensities (FIs) of CD34^+^CD7^+^CD4^+^ cells were compared between HIV^+^ and HIV--samples using the Mann-Whitney test. (C) Correlation between the CD4 and CD34 FI of CD34^+^CD7^+^CD4^+^ cells in the HIV^+^ sample of ID 1 were analysed (n = 3 for HIV-; n = 7 for HIV^+^). (D) CD34^+^CD7^+^CD4^+^CXCR4^+^ cells were tested for correlations with the frequencies of p24^+^CD34^+^ (left) and p24^+^CD34^+^CD7^+^ (right) cells. (E) Comparison of CD3 FIs between HIV-infected and uninfected samples. (F) Correlation analysis between CD3 FIs and CD4 (left) or CD34 (right) FIs. Comparisons were made using the Wilcoxon matched-pairs signed rank test unless otherwise noted. Spearman’s correlation coefficients were calculated for correlation analyses.

**Extended Data Figure 9.**
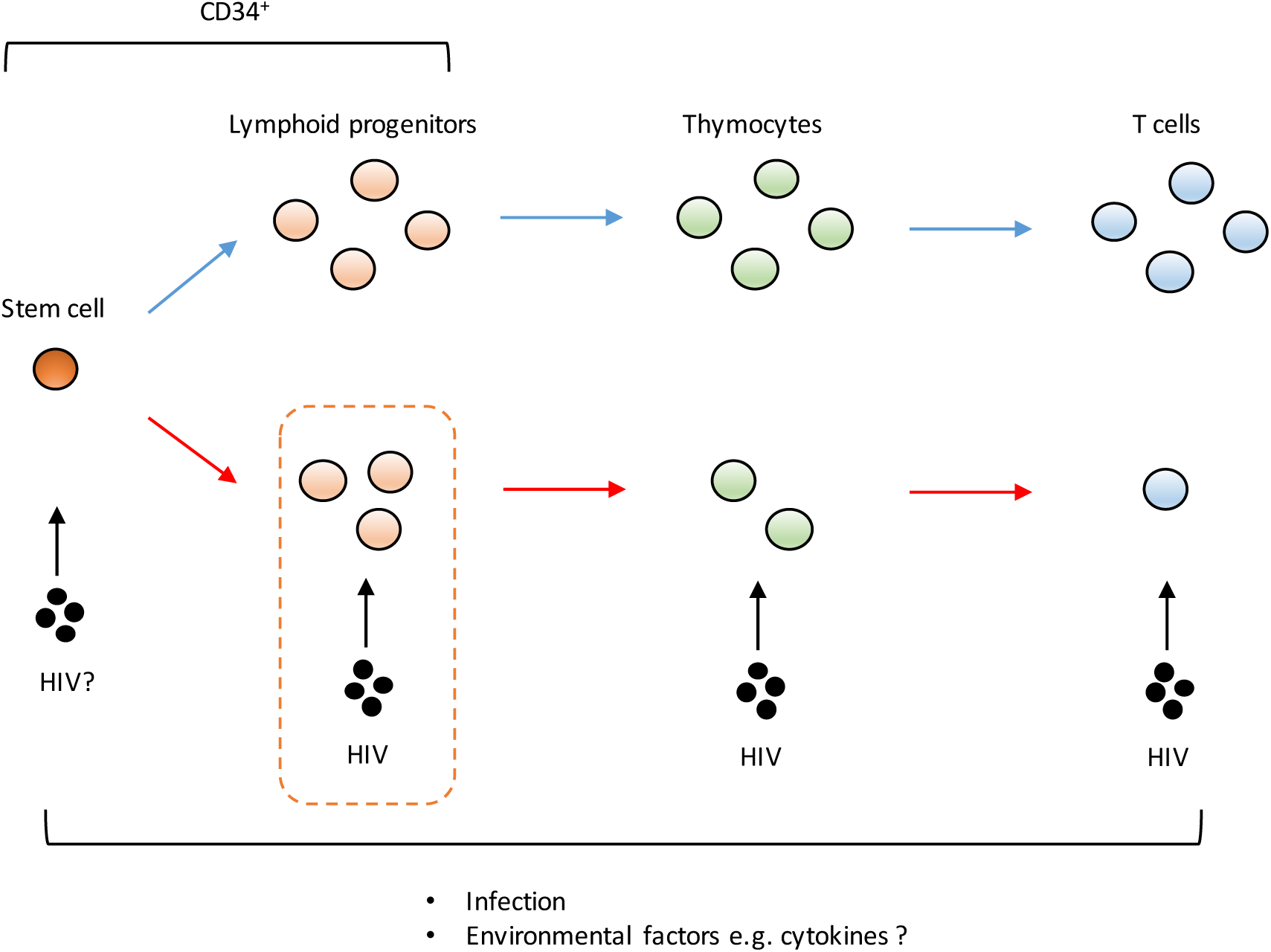
A schematic representation of factors potentially influencing the decrement of CD4^+^ T cells. Thymocytes and T cells are highly susceptible to HIV infection. However, the decrease of CD34^+^ lymphoid progenitor counts occurs early and precedes the decrease of thymocyte and T-cell counts. Although the mechanisms for the decrease of CD34^+^ lymphoid progenitors are unclear, this decline may limit the production of thymocytes and T cells.

**Extended Data Table 1.**
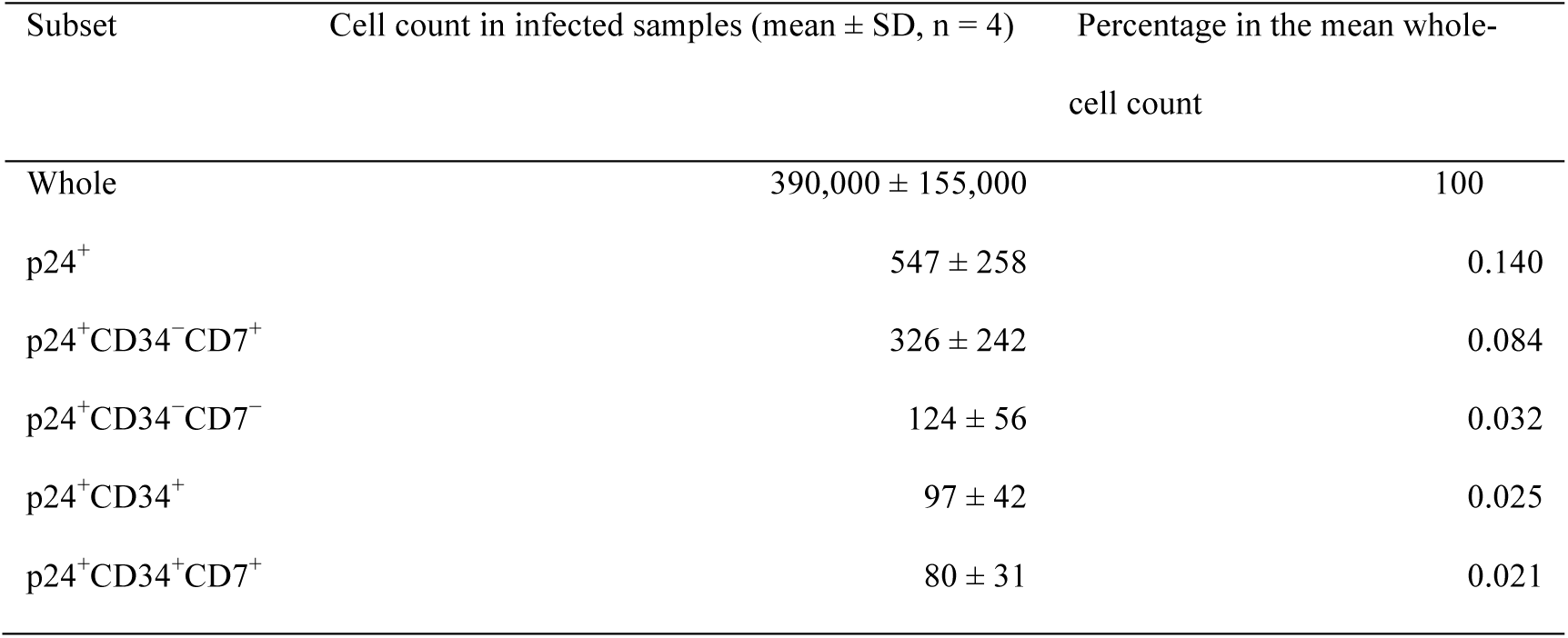
Mean cell counts (n = 4) in different HIV p24^+^ subsets of OP9-DL1 cocultured cells. The whole-cell count data were obtained using four of nine samples from experiment 2 (short coculture), in which cord-derived CD34^+^ cells were first cocultured with OP9-DL1 cells for 4–6 weeks, infected with HIV-1NL4-3, cocultured again for 1 week and analyzed by flow cytometry. The percentages in the mean whole-cell count are also shown, and they are comparable to those presented in Fig. 4C. SD, standard deviation.

| Subset | Cell count in infected samples (mean ± SD, n = 4) | Percentage in the mean whole-cell count |
| --- | --- | --- |
| Whole | 390,000 ± 155,000 | 100 |
| p24<sup>+</sup> | 547 ± 258 | 0.140 |
| p24<sup>+</sup>CD34<sup>-</sup>CD7<sup>+</sup> | 326 ± 242 | 0.084 |
| p24<sup>+</sup>CD34<sup>-</sup>CD7<sup>-</sup> | 124 ± 56 | 0.032 |
| p24<sup>+</sup>CD34<sup>+</sup> | 97 ± 42 | 0.025 |
| p24<sup>+</sup>CD34<sup>+</sup>CD7<sup>+</sup> | 80 ± 31 | 0.021 |

**Extended Data Table 2.**
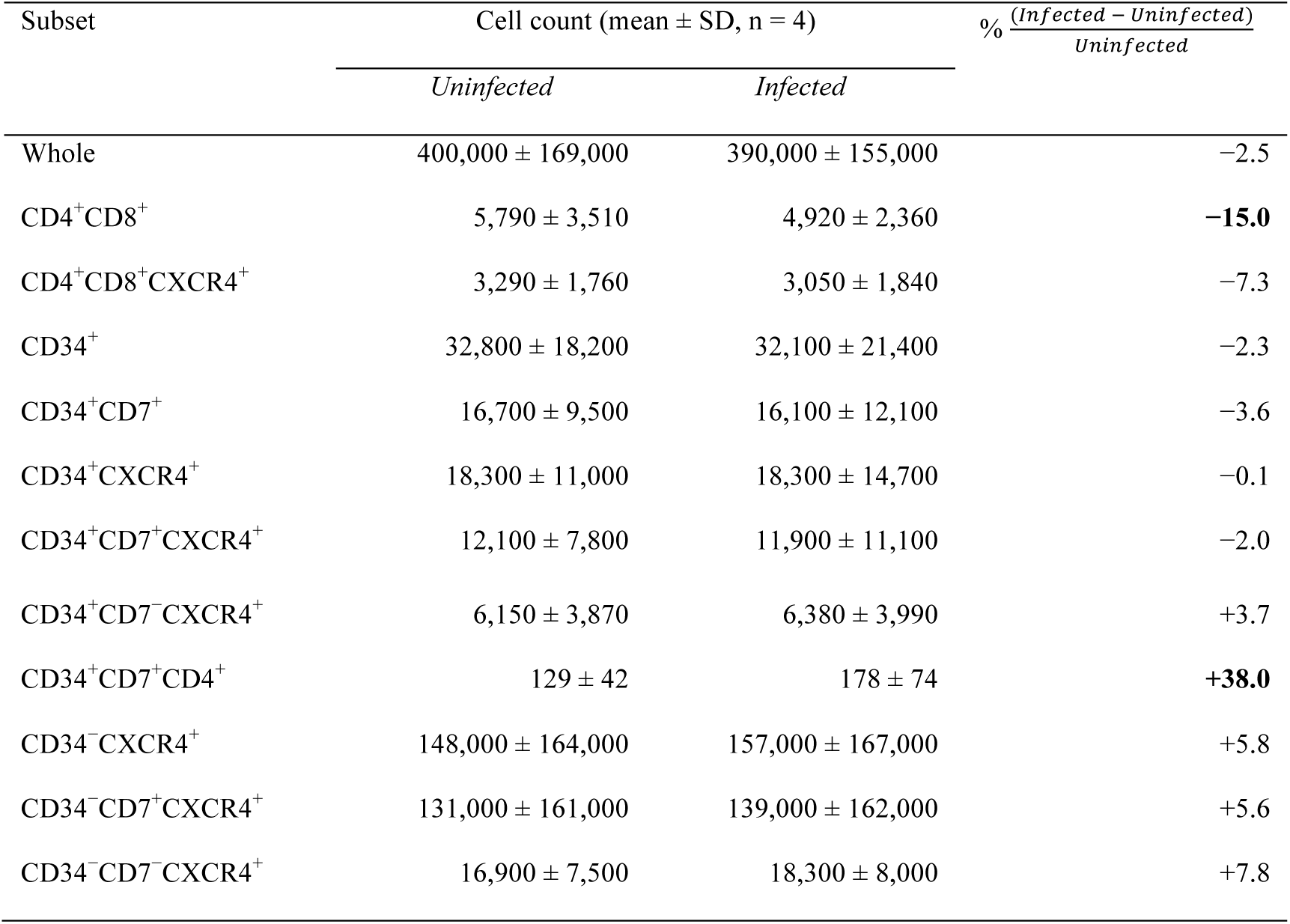
Mean cell counts (n = 4) in different subsets of OP9-DL1– cocultured cells. The whole-cell count data were obtained using four of nine samples from experiment 2 (short coculture), in which cord-derived CD34^+^ cells were first cocultured with OP9-DL1 for 4–6 weeks, infected with HIV-1NL4-3, cocultured again for 1 week and analyzed by flow cytometry. Cell counts in HIV-infected samples were compared with those in uninfected control samples. Percent changes in the subset cell counts following HIV infection were calculated, as shown in the right column. Percent changes of >10% are shown in bold. SD, standard deviation.

## References

1 Parinitha, S. & Kulkarni, M. Haematological changes in HIV infection with correlation to CD4 cell count. Australas Med J 5, 157–162, doi:10.4066/AMJ.20121008 (2012)

2 Corbeau, P. & Reynes, J. Immune reconstitution under antiretroviral therapy: the new challenge in HIV-1 infection. Blood 117, 5582–5590, doi:10.1182/blood-2010-12-322453 (2011)

3 Blom, M., Epeldegui, M. & Uittenbogaart, C. H. in HIV-Host Interactions (ed Theresa Li-Yun Chang) (InTech, 2011).

4 Ye, P., Kirschner, D. E. & Kourtis, A. P. The thymus during HIV disease: role in pathogenesis and in immune recovery. Curr HIV Res 2, 177–183 (2004)

5 Knutsen, A. P., Roodman, S. T., Freeman, J. J., Mueller, K. R. & Bouhasin, J. D. Inhibition of thymopoiesis of CD34^+^ cell maturation by HIV-1 in an in vitro CD34^+^ cell and thymic epithelial organ culture model. Stem Cells 17, 327–338, doi:10.1002/stem.170327 (1999)

6 Alexaki, A. & Wigdahl, B. HIV-1 infection of bone marrow hematopoietic progenitor cells and their role in trafficking and viral dissemination. PLoS Pathog 4, e1000215, doi:10.1371/journal.ppat.1000215 (2008)

7 Dhurve, S. A. & Dhurve, A. S. Bone Marrow Abnormalities in HIV Disease. Mediterr J Hematol Infect Dis 5, e2013033, doi:10.4084/MJHID.2013.033 (2013)

8 Banda, N. K. et al. Depletion of CD34^+^ CD4^+^ cells in bone marrow from HIV-1-infected individuals. Biol Blood Marrow Transplant 5, 162–172, doi:10.1053/bbmt.1999.v5.pm10392962 (1999)

9 Nakano, T., Kodama, H. & Honjo, T. Generation of lymphohematopoietic cells from embryonic stem cells in culture. Science 265, 1098–1101 (1994)

10 Schmitt, T. M. et al. Induction of T cell development and establishment of T cell competence from embryonic stem cells differentiated in vitro. Nat Immunol 5, 410–417, doi:10.1038/ni1055 (2004)

11 De Smedt, M., Hoebeke, I. & Plum, J. Human bone marrow CD34^+^ progenitor cells mature to T cells on OP9-DL1 stromal cell line without thymus microenvironment. Blood Cells Mol Dis 33, 227–232, doi:10.1016/j.bcmd.2004.08.007 (2004)

12 Janas, M. L. et al. Thymic development beyond beta-selection requires phosphatidylinositol 3-kinase activation by CXCR4. J Exp Med 207, 247–261, doi:10.1084/jem.20091430 (2010)

13 Mohtashami, M. et al. Direct comparison of Dll1- and Dll4-mediated Notch activation levels shows differential lymphomyeloid lineage commitment outcomes. J Immunol 185, 867–876, doi:10.4049/jimmunol.1000782 (2010)

14 Weiss, R. A. HIV receptors and the pathogenesis of AIDS. Science 272, 1885–1886 (1996)

15 Grivel, J. C. & Margolis, L. B. CCR5- and CXCR4-tropic HIV-1 are equally cytopathic for their T-cell targets in human lymphoid tissue. Nat Med 5, 344–346, doi:10.1038/6565 (1999)

16 Schnittman, S. M. et al. Preferential infection of CD4^+^ memory T cells by human immunodeficiency virus type 1: evidence for a role in the selective T-cell functional defects observed in infected individuals. Proc Natl Acad Sci U S A 87, 6058–6062 (1990)

17 Okoye, A. A. & Picker, L. J. CD4(+) T-cell depletion in HIV infection: mechanisms of immunological failure. Immunol Rev 254, 54–64, doi:10.1111/imr.12066 (2013)

18 Kucia, M. et al. CXCR4-SDF-1 signalling, locomotion, chemotaxis and adhesion. J Mol Histol 35, 233–245 (2004)

19 Lapidot, T. & Kollet, O. The essential roles of the chemokine SDF-1 and its receptor CXCR4 in human stem cell homing and repopulation of transplanted immune-deficient NOD/SCID and NOD/SCID/B2m(null) mice. Leukemia 16, 1992–2003, doi:10.1038/sj.leu.2402684 (2002)

20 Kucia, M. et al. Trafficking of normal stem cells and metastasis of cancer stem cells involve similar mechanisms: pivotal role of the SDF-1-CXCR4 axis. Stem Cells 23, 879–894, doi:10.1634/stemcells.2004-0342 (2005)

21 Plotkin, J., Prockop, S. E., Lepique, A. & Petrie, H. T. Critical role for CXCR4 signaling in progenitor localization and T cell differentiation in the postnatal thymus. J Immunol 171, 4521–4527 (2003)

22 Berkowitz, R. D., Beckerman, K. P., Schall, T. J. & McCune, J. M. CXCR4 and CCR5 expression delineates targets for HIV-1 disruption of T cell differentiation. J Immunol 161, 3702–3710 (1998)

23 Suzuki, H. et al. Intrathymic effect of acute pathogenic SHIV infection on T-lineage cells in newborn macaques. Microbiol Immunol 49, 667–679, doi:10.1111/j.1348-0421.2005.tb03646.x (2005)

24 Marsden, M. D. et al. HIV latency in the humanized BLT mouse. J Virol 86, 339–347, doi:10.1128/jvi.06366-11 (2012)

25 Dudek, T. E. & Allen, T. M. HIV-specific CD8(+) T-cell immunity in humanized bone marrow-liver-thymus mice. J Infect Dis 208 Suppl 2, S150–154, doi:10.1093/infdis/jit320 (2013)

26 Pace, M. & O’Doherty, U. Hematopoietic stem cells and HIV infection. J Infect Dis 207, 1790–1792, doi:10.1093/infdis/jit120 (2013)

27 Hunt, P. W. et al. T cell activation is associated with lower CD4^+^ T cell gains in human immunodeficiency virus-infected patients with sustained viral suppression during antiretroviral therapy. J Infect Dis 187, 1534–1543, doi:10.1086/374786 (2003)

28 Isgro, A. et al. Altered clonogenic capability and stromal cell function characterize bone marrow of HIV-infected subjects with low CD4^+^ T cell counts despite viral suppression during HAART. Clin Infect Dis 46, 1902–1910, doi:10.1086/588480 (2008)

29 Shirozu, M. et al. Structure and chromosomal localization of the human stromal cell-derived factor 1 (SDF1) gene. Genomics 28, 495–500, doi:10.1006/geno.1995.1180 (1995)

30 Moll, N. M. & Ransohoff, R. M. CXCL12 and CXCR4 in bone marrow physiology. Expert Rev Hematol 3, 315–322, doi:10.1586/ehm.10.16 (2010)

31 Petrie, H. T. Cell migration and the control of post-natal T-cell lymphopoiesis in the thymus. Nat Rev Immunol 3, 859–866, doi:10.1038/nri1223 (2003)

32 Nixon, C. C. et al. HIV-1 infection of hematopoietic progenitor cells in vivo in humanized mice. Blood 122, 2195–2204, doi:10.1182/blood-2013-04-496950 (2013)

33 Carter, C. C. et al. HIV-1 utilizes the CXCR4 chemokine receptor to infect multipotent hematopoietic stem and progenitor cells. Cell Host Microbe 9, 223–234, doi:10.1016/j.chom.2011.02.005 (2011)

34 Ho Tsong Fang, R., Colantonio, A. D. & Uittenbogaart, C. H. The role of the thymus in HIV infection: a 10 year perspective. AIDS 22, 171–184, doi:10.1097/QAD.0b013e3282f2589b (2008)

35 Bordoni, V. et al. Early ART in primary HIV infection may also preserve lymphopoiesis capability in circulating haematopoietic progenitor cells: a case report. J Antimicrob Chemother, doi:10.1093/jac/dku559 (2015)

36 Akkina, R. New insights into HIV impact on hematopoiesis. Blood 122, 2144–2146, doi:10.1182/blood-2013-08-518274 (2013)

37 Dorival, C. et al. HIV-1 Nef protein expression in human CD34^+^ progenitors impairs the differentiation of an early T/NK cell precursor. Virology 377, 207–215, doi:10.1016/j.virol.2008.04.009 (2008)

38 Hoebeke, I. et al. T-, B- and NK-lymphoid, but not myeloid cells arise from human CD34(+)CD38(-)CD7(+) common lymphoid progenitors expressing lymphoid-specific genes. Leukemia 21, 311–319, doi:10.1038/sj.leu.2404488 (2007)

39 Awong, G. et al. Characterization in vitro and engraftment potential in vivo of human progenitor T cells generated from hematopoietic stem cells. Blood 114, 972–982, doi:10.1182/blood-2008-10-187013 (2009)

40 Bordoni, V. et al. Bone Marrow CD34^+^ Progenitor Cells from HIV-Infected Patients Show an Impaired T Cell Differentiation Potential Related to Proinflammatory Cytokines. AIDS Res Hum Retroviruses 33, 590–596, doi:10.1089/aid.2016.0195 (2017)

41 Savkovic, B. et al. A quantitative comparison of anti-HIV gene therapy delivered to hematopoietic stem cells versus CD4^+^ T cells. PLoS Comput Biol 10, e1003681, doi:10.1371/journal.pcbi.1003681 (2014)

42 Liu, Y., Zhou, J., Pan, J. A., Mabiala, P. & Guo, D. A novel approach to block HIV-1 coreceptor CXCR4 in non-toxic manner. Mol Biotechnol 56, 890–902, doi:10.1007/s12033-014-9768-7 (2014)

43 Suzuki, K., Ahlenstiel, C., Marks, K. & Kelleher, A. D. Promoter Targeting RNAs: Unexpected Contributors to the Control of HIV-1 Transcription. Mol Ther Nucleic Acids 4, e222, doi:10.1038/mtna.2014.67 (2015)

44 Costin, J. M. Cytopathic mechanisms of HIV-1. Virol J 4, 100, doi:10.1186/1743-422x-4-100 (2007)

45 Tsukamoto, T., Yamamoto, H., Okada, S. & Matano, T. Recursion-based depletion of human immunodeficiency virus-specific naive CD4(+) T cells may facilitate persistent viral replication and chronic viraemia leading to acquired immunodeficiency syndrome. Med Hypotheses 94, 81–85, doi:10.1016/j.mehy.2016.06.024 (2016)

46 Su, L. et al. HIV-1-induced thymocyte depletion is associated with indirect cytopathogenicity and infection of progenitor cells in vivo. Immunity 2, 25–36 (1995)

47 Adachi, A. et al. Production of acquired immunodeficiency syndrome-associated retrovirus in human and nonhuman cells transfected with an infectious molecular clone. J Virol 59, 284–291 (1986)

48 Holmes, R. & Zuniga-Pflucker, J. C. The OP9-DL1 system: generation of T-lymphocytes from embryonic or hematopoietic stem cells in vitro. Cold Spring Harb Protoc 2009, pdb.prot5156, doi:10.1101/pdb.prot5156 (2009)

49 Tsukamoto, T. & Okada, S. The use of RetroNectin in studies requiring in vitro HIV-1 infection of human hematopoietic stem/progenitor cells. J Virol Methods 248, 234–237, doi:10.1016/j.jviromet.2017.08.003 (2017)

50 Suzuki, K. et al. Prolonged transcriptional silencing and CpG methylation induced by siRNAs targeted to the HIV-1 promoter region. J RNAi Gene Silencing 1, 66–78 (2005)

